# A Multi-Scale Study of Thalamic State-Dependent Responsiveness

**DOI:** 10.1101/2023.12.02.567941

**Authors:** Jorin Overwiening, Federico Tesler, Domenico Guarino, Alain Destexhe

## Abstract

The thalamus is the brain’s central relay station, orchestrating sensory processing and cognitive functions. However, how thalamic function depends on internal and external states, is not well understood. A comprehensive understanding would necessitate the integration of single cell dynamics with their collective behavior at population level. For this we propose a biologically realistic mean-field model of the thalamus, describing thalamocortical relay neurons (TC) and thalamic reticular neurons (RE). We perform a multi-scale study of thalamic responsiveness and its dependence on cell and brain states. Building upon existing single-cell experiments we show that: (1) Awake and sleep-like states can be defined via the absence/presence of the neuromodulator acetylcholine (ACh), which controls bursting in TC and RE. (2) Thalamic response to sensory stimuli is linear in awake state and becomes nonlinear in sleep state, while cortical input generates nonlinear response in both awake and sleep state. (3) Stimulus response is controlled by cortical input, which suppresses responsiveness in awake state while it ‘wakes-up’ the thalamus in sleep state promoting a linear response. (4) Synaptic noise induces a global linear responsiveness, diminishing the difference in response between thalamic states. Finally, the model replicates spindle oscillations within a sleep-like state, exhibiting a qualitative change in activity and responsiveness. The development of this novel thalamic mean-field model provides a new tool for incorporating detailed thalamic dynamics in large scale brain simulations.

**Author summary:** The thalamus is a fascinating brain region that acts as the gate for information flow between the brain and the external world. While its role and importance in sensory and motor functions is well-established, recent studies suggest it also plays a key role in higher-order functions such as attention, sleep, memory, and cognition. However, understanding how the thalamus acts on all these functions is challenging due to its complex interactions at both the neuron level and within larger brain networks. In this study, we used a mathematical model grounded in experimental data that realistically captures the behavior of the thalamus, connecting the scales of individual neurons with larger populations. We found that the thalamus functions differently depending on whether the brain is in an awake or a sleep-like state: When awake, the thalamus processes sensory information in a straightforward way, resulting in a faithful information transmission to the cortex. But during sleep, only significant or important stimuli create a response. Importantly, this behavior can be controlled by cortical-like input and noise. With this study, we shed light on how the thalamus might modulate and interact with various brain functions across different scales and states. This research provides a deeper understanding of the thalamus’s role and could inform future studies on sleep, attention, and related brain disorders.

## 1 Introduction

The thalamus, a well preserved structure found in all mammals [1], serves as the core relay hub of the central nervous system. Diverse thalamic nuclei function as transmitters of sensory information from the periphery to the cortex and other central nervous system structures, while also facilitating the transfer of motor commands from the cortex to various regions of the body [2, p. 4-5]. Each of the relatively independent thalamic nuclei comprises at least two cell types: excitatory (glutamergic) principal *relay* cells, featuring extensive axonal projections to various nervous system structures, but rarely to other principal cells, and local inhibitory (GABAergic) interneurons [3, 4].

The primary source of activity in thalamic nuclei arises from direct pathways, operating in both peripheral-to-central and central-to-peripheral directions. Additionally, cortical feedback projections exert a strong influence on the thalamus. Notably, the number of thalamo-cortical outgoing axons is approximately one-tenth of the number of cortico-thalamic incoming axons [3, 5], and the cortex is the major source of synapses within the thalamus, for example accounting for 50% of synapses in the lateral geniculate nucleus (LGN) [3]. This extensive feedback loop between the thalamus and cortex indicates a substantial modulating role of the cortex in thalamic relay functions [6].

During attentive wakefulness, thalamic relay neurons display tonic firing. However, membrane hyperpolarization leads to bursting behavior via low-threshold Ca^+^ channels [7]. Bursting occurs in deep sleep states (NREM) and general states of low attention [8], in which hyperpolarization is generated by a low level of the neuromodulator acetylcholine (ACh) [9].

Surrounding the thalamus, the thalamic reticular nucleus (TRN) contains GABAergic reticular cells (RE) that broadly inhibit thalamic nuclei through axonal, and themselves through dense axonal and dendritic connections [3, 7]. RE neurons can be activated through feedforward signals from thalamic nuclei or feedback from the cortex. These neurons consistently exhibit bursting behavior and can induce similar patterns in thalamic relay cells via hyperpolarization. This recurrent network allows the cortex and thalamus itself to actively modulate thalamic response and transfer of information, rendering the thalamus as a gate, with the TRN as the gatekeeper.

In addition to its gating function, there is also evidence of the thalamus playing a principal role in whole brain dynamics, such as spindle oscillations or slow waves in NREM sleep or anaesthesia [10–12]. It is suggested that the intrinsic loop between thalamocortical (TC) relay and RE cells plays a pivotal role in all of these behaviours by acting as a pacemaker and oscillator. The crucial mechanism at play is the rebound bursting of relay cells via hyperpolarization induced by RE inhibition.

These oscillatory behaviors primarily manifest during sleep-like brain states, where the TC cells show a prevalence for bursting [13]. Additionally, in newer studies it was shown that thalamic integration with cortical pathways suggests a significant role of the thalamus in many higher brain functions, including sensation, attention, and cognition [14, 15].

Investigating the interaction between thalamic reticular and relay neurons at various levels is therefore crucial for deciphering the interplay of the brain with the outside world. To this end it is necessary to analyse these neuron interactions and their corresponding population activity via large-scale models. One feasible approach for scaling upwards is to employ networks of single-cell neuron models, but the computational demand rapidly increases as the network size is taken to the scale of anatomical subdivisions of the brain. For larger scales and even whole-brain simulations, it is necessary to decrease computational complexity. This can be achieved by reducing the degrees of freedom and describing homogeneous populations of neurons as the smallest units. A viable option is to use a mean-field theory to model population dynamic statistics.

Most existing neuronal field models can be separated in two groups: either phenomenological models (e.g. [16–18]), or more abstract mathematical models (e.g. [19–21]). Phenomenological models replicate biological behaviour and are capable of modelling particular brain regions, cell types or whole brain recordings. However, these can not couple significant effects or characteristics to model parameters which makes it impossible to use such models far of the fitting point and renders analytical analysis impractical. Conversely, abstract mathematical models couple the dynamical aspects of neuronal activity directly to model parameters and allow analytical or fast-forward numerical analysis, but model parameters are often not well linked to biological observables.

To strike a good balance between these two options, we develop in this paper a *biologically realistic* mean-field model of the thalamus that also allows analytical analysis. To achieve this biological realism with a firing rate model, our formalism follows a bottom-up approach, starting at the single-cell level and incorporating cellular and structural specificities of the thalamic circuits [22]. Our approach incorporates three crucial biological features: (1) *Irregular spiking* activity of neurons is believed to be important for transfer efficiency [23] and the correct baseline for neurons in both awake-like asynchronous (AI) states [24] as well as in sleep-like synchronous (SI) states [25]. (2) Synaptic *conductances* allow for realistic bi-stability and self-sustained activity [26] as well as modeling the fluctuation-driven regime [27]. (3) *Adaptation* mechanisms are the main generators of the different firing behaviors in the brain and important to include into models for generating realistic firing rate saturation and especially the bursting behavior of thalamic cells.

Using this mean-field model we investigate the state-dependent responsiveness of the thalamus, integrating the interplay between multiple scales (from single-cell level to the mesoscale). Building upon existing single-cell experiments we show that: First, the transition from tonic to burst firing of TC cells via ACh renders thalamic response nonlinear in sleep state (Section 3.2). Second, sensory stimuli generate a linear response, while cortical inputs generate a nonlinear response of the thalamus (Section 3.3). Third, cortical input and synaptic noise modulate thalamic response and synaptic noise diffuses thalamic state transitions and removes thalamic response dependency on both voltage and frequency (Section 3.4). Finally, we demonstrate that the proposed model is capable of generating self-sustained spindle oscillations, drastically altering responsiveness in this state (Section 3.5).

## 2 Methods

In this section we describe the single-cell, network, and the mean-field model. The chosen network and connectivity structure as well as cell and synaptic parameters are described.

### 2.1 Spiking neuron model

For both single-cell and network simulations we employ the *Adaptive exponential integrate and fire model* (AdEx) (as defined in [28] and analysed in [29]). This conductance based model often proved to be a good balance between computability and biological realism in terms of capturing all firing modes observable in real cells [30] and significantly in thalamocortical cells [31, 32]. Importantly, it allows for a systematic fit of real cell traces. The dynamical system is the two equations describing membrane potential *v* and adaptation current *ω* of a given cell *µ*

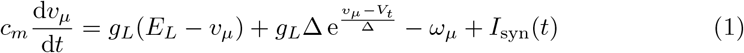

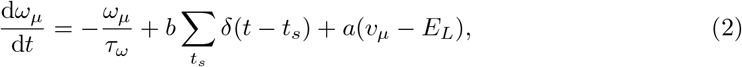

with the cell parameters listed in Table 1 and where *I*_syn_ models all incoming synaptic currents. It consists of two currents dependent on excitatory 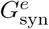 and inhibitory 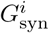 membrane conductances and is defined as

**Table 1.** Cell and synaptic parameters for TC and RE cells in awake (ACh) and sleep (no ACh) states. Connection parameters see Fig. 1a. The last two parameters are for the spiking network only and ”-” means the same value as in awake state.

| PARAM | AWAKE |  | SLEEP |  | DESCRIPTION |
| --- | --- | --- | --- | --- | --- |
|  | TC | RE | TC | RE |  |
| $Q_e$ | 1nS | 4nS | - | - | exc.quant. conductance increment |
| $Q_i$ | 6nS | 1nS | - | - | inh.quant. conductance increment |
| $c_m$ | 160pF | 200pF | - | - | membrane capacitance |
| $E_L$ | -65mV | -75mV | -70mV | -85mV | resting (leakage) potential |
| $g_L$ | 10nS | 10nS | 9.5nS | 13nS | leak conductance |
| $\tau_w$ | 200ms | 200ms | 270ms | 230ms | adaptation time const. |
| $a$ | 0nS | 8nS | 24nS | 28nS | membrane potential adaptation |
| $b$ | 10pA | 10pA | 200pA | 20pA | spike frequency adaptation |
| $\tau_{(e,i)}$ | 5ms | 5ms | - | - | syn. time constants |
| $E_e$ | 0mV | 0mV | - | - | exc. reversal potential |
| $E_i$ | -80mV | -80mV | - | - | inh. reversal potential |
| $V_t$ | -50mV | -45mV | - | - | threshold for the spike onset |
| $\Delta$ | 4.5mV | 2.5mV | - | - | amplitude of the spike onset |

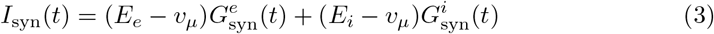

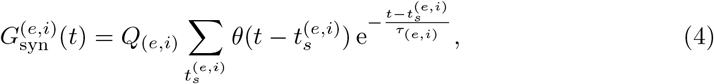

where *G*_syn_ is modeled such that each time a spike (*t*_*s*_) arrives these conductances experience an increment *Q* and exponentially relax again with time constant *τ*. As a baseline we use *Q*_*e*_ = 1nS and *Q*_*i*_ = 5nS [22].

Additional to the integration of this ODE set comes the usual spike mechanism employed in integrate and fire models: A spike of neuron *µ* is counted if *v*_*µ*_ *>V*_thr_ = −20mV, then the membrane potential is reset to *V*_*r*_ = {−55mV for RE, −50mV for TC} for a refractory period of 5ms.

### 2.2 Network architecture and model parameters

We model one thalamocortical relay (TC) and one connected reticular (RE) population of a generic lateral thalamic nucleus. We neglect interneurons, as it can be assumed that they only yield minor contribution to population dynamics [33]. One of the main potential application of the thalamus mean-field is to be incorporated into large or whole brain models with already developed cortical and sub-cortical mean-field models and related implementations [22, 34–40]. As a reference, these previous works on cortical circuits describe typically populations of ∼10^4^ neurons, corresponding to the size of a single cortical column. To keep the scale difference between cortex and thalamus proportional, we employ a scale of 1*/*10 [3, 5, 41] and therefore use *N* = 500 neurons per population. This allows to build a basic realistic-scale thalamo-cortical loop with just two mean-field models.

The network with its connections is depicted in Fig. 1a. We consider a random connected *Erdos-Renyi* network comparable to the statistical assumptions of the meanfield (Table S1). TC and RE populations form a loop of excitation and inhibition. TC cells do not excite other TC cells but RE cells (next to outgoing axons to the cortex). In contrast, RE are connected in an inhibitory loop and also inhibit TC cells. We propose two external drives serving as inputs to the model: The *cortical* drive *P* (going to both populations) and the *sensory* drive *S* (going only to TC cells) modeling cortical signals and sensory stimuli to the thalamus, respectively.

**Fig. 1.**
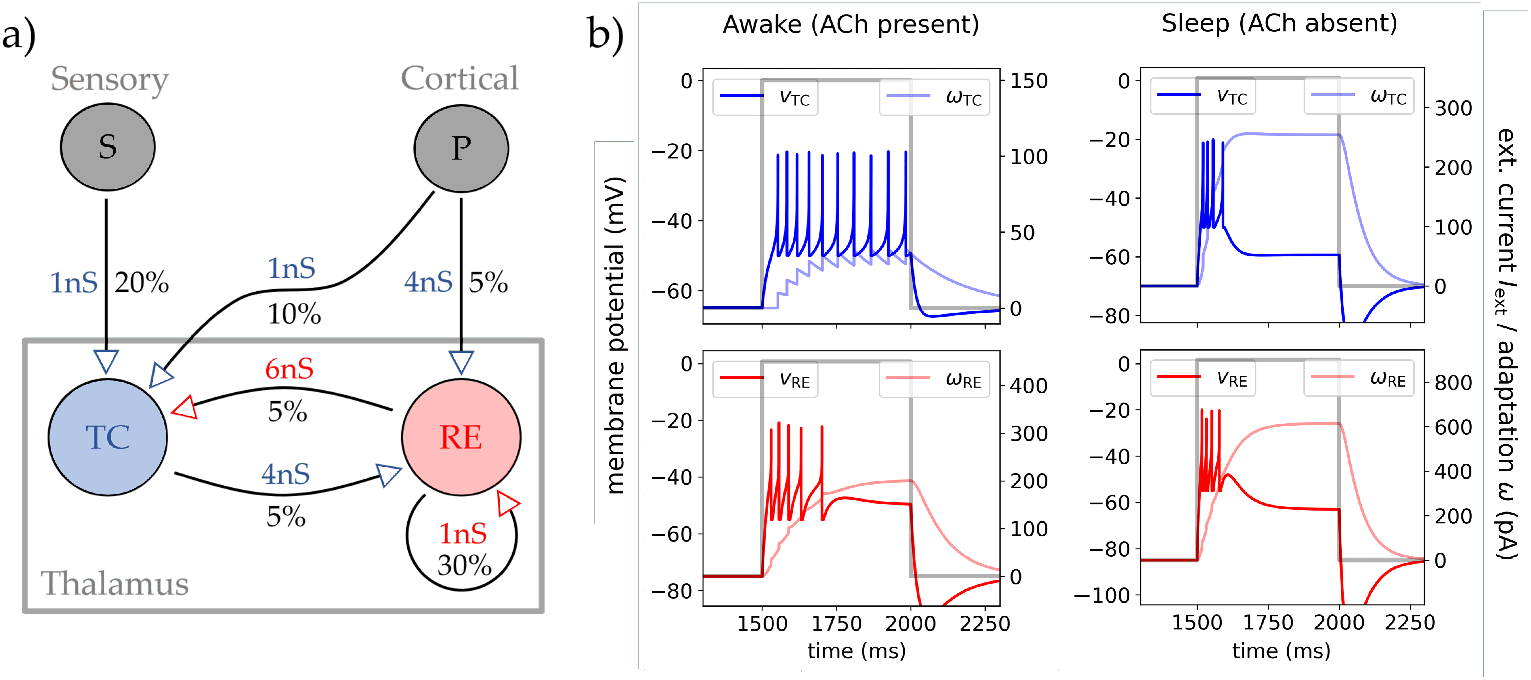
Network structure and single cell dynamics. **(a)** The chosen network structure of two thalamic populations (TC and RE cells of each *N* = 500), their synaptic increments *Q*, and connection probabilities *p* for all connection between the TC and RE population. The external inputs are shown in grey: The *cortical* drive *P* (*N* = 8000) and the *sensory* stimulus drive *S* (*N* = 500). The arrows mark the direction of synaptic transmission and if they act excitatory (blue) or inhibitory (red). **(b)** Single cell traces from AdEx IF neurons (see Methods for details) of TC and RE for a timed constant input current (grey line). The left column shows TC and RE response to the injected current in awake state (with ACh) and the right column the same in sleep state (no ACh). The cells membrane potential *v* and adaptation current *ω* are shown in color for TC (blue) and RE cell (red), respectively.

For the synaptic and connection parameter values, we start with a connection probability between TC and RE populations of *p* = 5%, which captures the sparse connectivity between the two populations [4]. To model the dense net of locally self-inhibiting RE neurons in the TRN [2, 42, 43], we use *p* = 30%. There are 2 to 10 times more axons projecting from cortex to TC than from cortex to RE cells, but the amplitude of connection to RE is stronger, keeping a strong inhibitory corticothalamic modulation via the TRN [3, 7]. Last, the number and convergence of axons from RE to TC cells ensures sparse but strong inhibition [4, 7]. See Fig. 1a for all the parameter values.

Moving to cell parameters, we model two states of the thalamus corresponding to high or low levels of the excitatory modulator acetylcholine (ACh). In McCormick and Prince [9] and [44], it was shown that low levels of ACh change the firing patterns of TC cells to inhibit single tonic firing and to promote bursting. Because of the capability of ACh to act as a switch between tonic and bursting mode in the TC cells relay, and its role in controlling the overall physiological brain state [33, 45, 46], we define here these two states as *awake* state (ACh present; wakefulness, REM sleep) and *sleep* state (ACh absent; NREM sleep, low attention).

Parameters of the single-cell model were determined via Mean-Absolute-Error (MAE) analysis between model prediction and recorded TC and RE cell traces of those two studies [9, 44] for the two states of ACh present (awake) and ACh absent (sleep). Initial parameter values are taken from Destexhe [31]. Parameter ranges were constrained by experimentally observed ranges extracted from [9, 44] and the Neuro-Electro database [47] for TC and RE neurons. The resulting parameters are shown in Table 1. The robustness of the model parameters is validated via a large range exploration in the parameter space, shown in Fig. S4–S7. Beyond this initial determination, two further modifications to the parameters values were performed: (1) A hyperpolarised *E*_*L*_ for RE cells in sleep state. This choice does deviate from a best fit as is evident from the increased error as shown in the second row in Fig. S6. However, this was necessary to guarantee biologically realistic stable and balanced AI dynamics of the full network (see Section 3.1). And (2) in a stronger spike adaptation *b* for TC cells, also in sleep state. This choice is required for TC cells to burst also in network simulations (see Fig. 3.b) and leads to stronger bursting at the single-cell level, but does not significantly increase the error of the single-cell fit.

To show that the cells inherit the correct behaviour, using the AdEx (1) with the proposed cell parameters, in Fig. 1b four exemplary single cell traces of RE and TC are shown. In there a constant–time gated–current was injected in to the cell to invoke a firing response of the cell. This was done by setting a rectangular pulse as *I*_syn_ in (1) (*I*_syn_ generates tonic and burst firing via two different bifurcations depending on excitability state, see [29]). The top row shows the wanted response types for the TC cell: Tonic firing with awake parameters (modulating ACh), and burst firing with sleep parameters (low-level of ACh). In the bottom row, RE cell’s respond via burst firing in both parameter states, but the burst duration decreases in sleep state while keeping the same amount of spikes (increased *burstiness*).

### 2.3 Mean-field model

El Boustani and Destexhe [48] developed a second-order mean-field formalism of differential equations describing the firing rate statistical moments of spiking networks. This general framework closes the statistical hierachy at second order and is applicable to any arbitrary neuron models as long as a characteristic transfer function can be defined. It is assumed that the network is a sparse and randomly connected *Erdos-Renyi model*. It is derived with the assumption of the system being in an E-I balanced AI state. This formalism is extended by including the slow dynamic effects of adaptation [22] so that the system is fully described by mean firing rate *ν*_*µ*_ and adaptation *ω*_*µ*_ for each neuron population *µ*. The differential equation system for this framework then reads

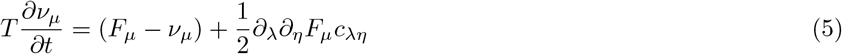

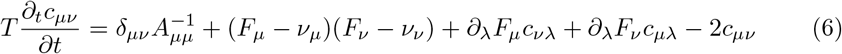

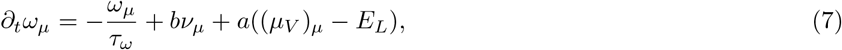

where *F*_*µ*_ is the transfer function of cell population *µ* and *c*_*µν*_ the covariance between two population’s firing activity. The indices {*µ, ν, λ, η*} run over the set of populations, e.g. in our case of two populations the set of {*e, i*} for excitatory TC and inhibitory RE. The derivatives are defined as 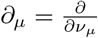. Important to note is the role of *T* which marks the adiabatic time step such that dynamics with smaller time resolutions are not captured and which has to fulfil the requirements of Table S1.

The core of this formalism is the transfer function *F* and so the main task in constructing a mean-field of the thalamus is to get the transfer function of TC and RE cells in the two states of awake and sleep.

To derive the transfer function we follow the semi-analytical approach of Zerlaut et al. [49] which combines the seminal studies of [50, 51]. In there the firing rate is written as a probabilistic function counting the spikes in term of the membrane potential *v*(*t*) being above a certain *spike threshold potential V*_*θ*_ in each time bin of duration *τ*_*V*_ which resembles the membrane potentials autocorrelation time. In the Gaussian limit we get a function dependent on the membrane subthreshold fluctuation statistical moments and define that as our transfer function

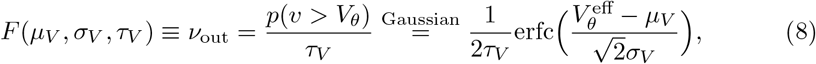

where *µ*_*v*_ is the mean and *σ*_*v*_ the standard deviation of the (subthreshold) membrane potential. In the second step, the constant threshold *V*_*θ*_ is replaced with a phenomenological one acting as a function dependent on – and therefore accounting for – different cell properties. Because there is no theoretical form, a general second order polynomial dependent on the set {*µ*_*V*_, *σ*_*V*_, *τ*_*V*_ } was proposed [51]

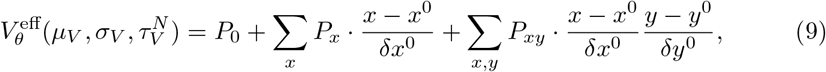

with 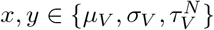 and where 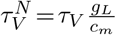 is the non-dimensionalised autocorrelation and the parameters space is normalised to limit the fluctuation driven regime, with mean *x*^0^ and deviation *δx*^0^.

Here either single cell simulations or experimental clamp data can be used to get values for the unknown amplitudes {*P*}. This fitting has to be performed for each distinctive cell type, so in our case for TC and RE neurons. Because the two states awake and sleep are mostly changes in adaptation parameters, and it was shown in [22] that the mean-field is predictive even far from its fitting point, we just need one fit per cell type. This also is biologically realistic, for the changes induced by e.g. ACh would not change the cell morphology, and we consider the threshold membrane potential to stay the same for both states.

The set of {*µ*_*V*_, *σ*_*V*_, *τ*_*V*_} can be calculated purely analytically by using Campbell’s theorem and assuming Poissonian distribution of incoming spikes as the generator of subthreshold fluctuations [50] (as is the case in the AI regime). The mean or static synaptic conductances are calculated then as a function of incoming spike frequencies {*ν*_*e*_, *ν*_*i*_} in terms of their mean and standard deviation

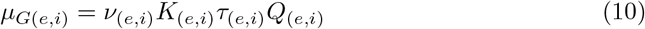

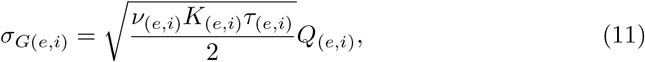

where *K*_*µ*_ = *p*_*µ*_*N*_*µ*_. With that the general input conductance of the cell can be computed

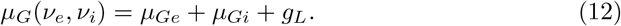

Then we can calculate the mean membrane potential *µ*_*V*_ from the first order approximation of (1) as a function of incoming spike frequencies

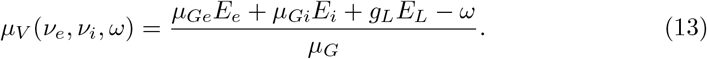

Taking (3) as the general synaptic input, we can calculate the form of a single postsynaptic potential (PSP). And via shotnoise theory get the density power spectrum of membrane fluctuations *P*_*V*_ (*q*) as a response to a stimulation (3). Then the variance of fluctuations with taking the integral in frequency domain 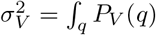, follows to

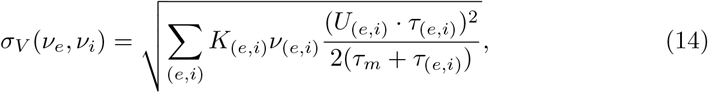

where 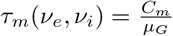 and 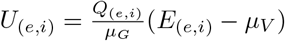 is the effective synaptic drive.

Finally, the autocorrelation time *τ*_*V*_ completes the framework which is defined in terms of the power spectrum as 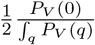, resulting in

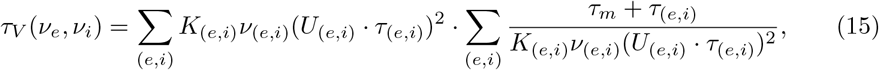

where in case of only one synaptic event this would reduce to *τ*_*V*_ = *τ*_*m*_ + *τ*_(*e,i*)_.

With (13), (14), and (15) the transfer function with effective threshold (8) is now dependent only on the incoming firing rates at excitatory and inhibitory synapses *F* (*µ*_*V*_, *σ*_*V*_, *τ*_*V*_) → *F* (*ν*_*e*_, *ν*_*i*_), closing our firing-rate based mean-field formalism.

### 2.4 Transfer function fit

To get the transfer functions we fit {*P*} on single cell simulations of TC and RE cells (in awake state) using the AdEx equations (1). The formalism translates excitatory and inhibitory input firing rates {*ν*_*e*_, *ν*_*i*_} of a neuron into its fluctuation statistics {*µ*_*V*_, *σ*_*V*_, *τ*_*V*_} and then to its output firing rate.

The advantage of this semi-analytic approach is that–given either simulated or experimental data–we can calculate the phenomenological threshold 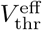 via reordering of (8). Then the employed procedure is to first fit (9) linearly in the threshold space (depending on the topography of the space to capture, this fit can be done nonlinearly too). However, here (13) has to be adjusted because the adaptation *ω* is unknown. Therefore, the (stationary) solution to (7) will be used to calculate *ω* from the firing rate data. The resulting values for *P* are following used as initial guesses for the fully nonlinear fit of (8) in the original firing rate space.

For the fit we normalised the fluctuation regime the same way as done in previous works [22, 51]; to ensure comparability: 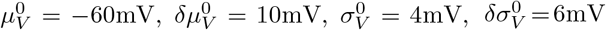, and 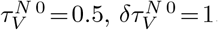.

## 3 Results

The results of this paper are structured as follow: First the mean-field model will be compared with simulated spiking network dynamics and validated in and far to from the fitting point (Section 3.1). Then thalamic responsiveness and how it depends on different external and internal states will be investigated (Bursting in Section 3.2, Inputs 3.3, and Noise 3.4). Lastly, spindle oscillations in a sleep-like state in the employed models are shown (Section 3.5).

### 3.1 Fitting and validation

In this section, we validate the mean-field model and demonstrate its suitability for modeling both awake and sleep state of the thalamus by comparing it with spiking networks.

The fit parameters of the mean-field’s transfer function via (9), obtained using our fitting technique (as described in Section 2.4), are depicted in Table 2. These parameters are applied to both awake and sleep states (ACh absent/present; see Section 2.2) and are used throughout this and the following three sections (but not in Section 3.5).

**Table 2.** The fitting parameters of 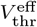 and their values. All values in mV.

| cell | $P_0$ | $P_\mu$ | $P_\sigma$ | $P_\tau$ | $P_{\mu\mu}$ | $P_{\mu\sigma}$ | $P_{\mu\tau}$ | $P_{\sigma\sigma}$ | $P_{\sigma\tau}$ | $P_{\tau\tau}$ |
| --- | --- | --- | --- | --- | --- | --- | --- | --- | --- | --- |
| TC | -47.31 | 1.68 | 0.97 | -3.46 | 0.47 | -1.68 | -6.46 | 3.43 | -1.14 | 0.19 |
| RE | -40.77 | -1.98 | -3.12 | 3.57 | 1.39 | -0.38 | -0.33 | 0.16 | 0.26 | -0.53 |

In Fig. 2a, we show the fitted transfer functions *F* for TC cells (top, blue) and RE cells (bottom, red) across the full range of excitatory input frequencies (*ν*_*e*_) and a subset of three inhibitory input frequencies (*ν*_*i*_). Each dot represents the averaged output frequency from the single-cell simulations over 5 seconds. The sigmoid shape of the transfer function (8) is evident. Certain deviations from the fitted predictions via *F* are observed only at very high firing rates; a region of lesser biological relevance for the phenomena studied in this paper. To improve statistics, the single-cell firing rates were averaged over 100 runs.

**Fig. 2.**
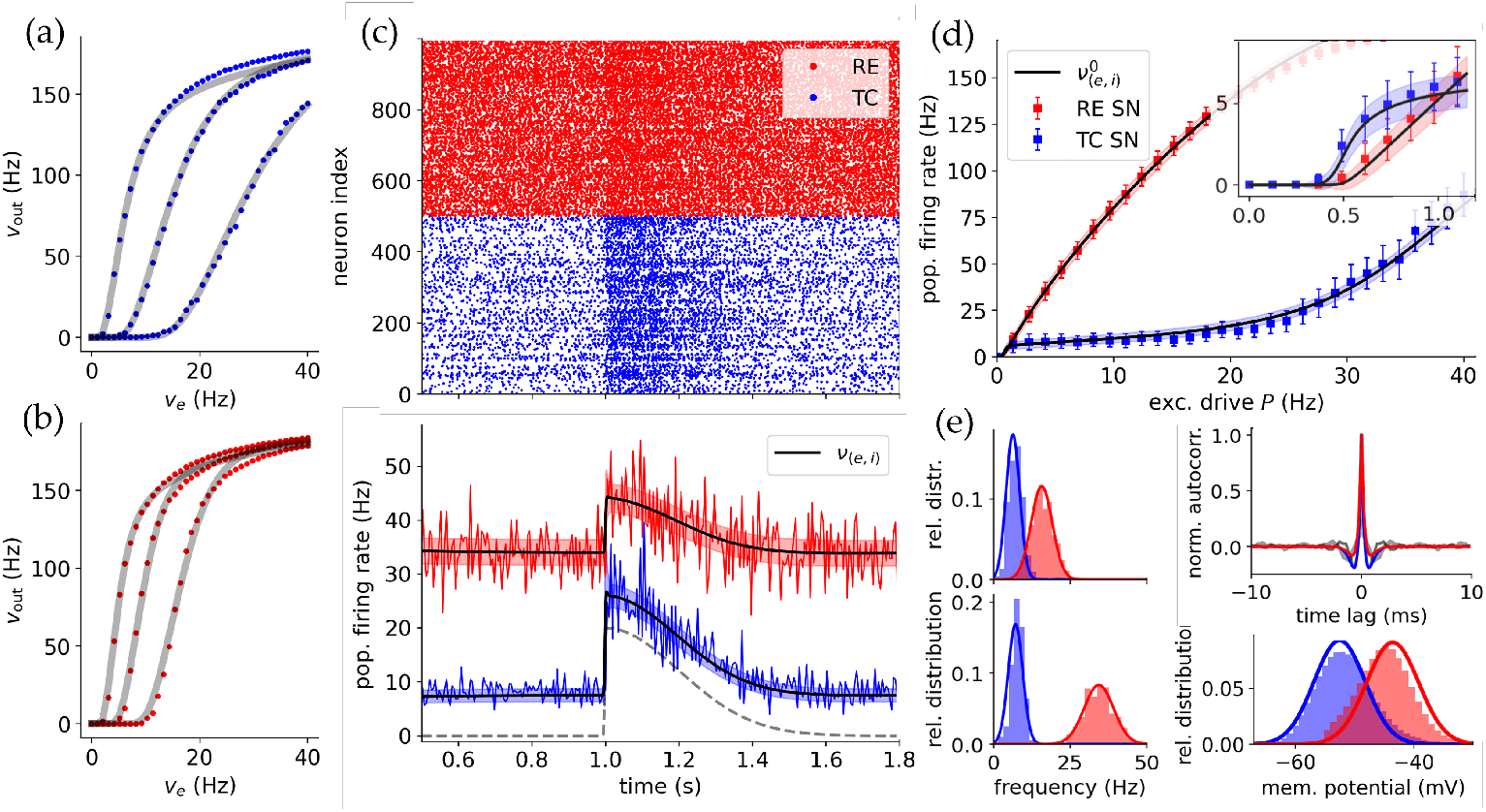
Validating the mean-field with spiking networks. **(a,b)** The fitted transfer functions for RE and TC cell for three different inhibitory inputs each with their corresponding single cell simulations. Top (a, blue) is for TC and bottom (b, red) for RE cell-type (in awake state). The dots each represent the averaged firing rate of a cell over 100 runs. **(c)** Comparison of the firing rate of the mean-field and the spiking network for constant cortical drive *P* = 4Hz and a split-Gaussian stimulus coming from *S*. Top is the raster plot showing all spiking times {*t*_*s*_} for all neurons in the spiking network simulation. Bottom is the averaged mean firing rate of spiking network (blue/red lines) and predicted mean firing rate of the mean-field *ν* (black line) with its standard deviation (shaded blue/red areas). **(d)** Comparison of the equilibrium firing rate of the spiking network and of the mean-field over a range of cortical inputs. Each dot represents a spiking network simulation for 10s where the steady long time mean is calculated. The black lines correspond to the mean-fields fixpoints 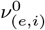, with the shaded areas being the standard deviations. The *inhibited* regime between ca. *P* ≃ (1, 20)Hz marks the standard activity employed. The inset shows a zoom at the low-drive regimes where activity is first silent and then controlled by TC until *P* ≃ 1Hz. **(e)** Left column: The firing rate distributions of spiking network (histogram) and mean-field (line) for *P* = {2, 4}Hz. Bottom-right: Comparison of membrane potential distribution for 4Hz. Top-right: Autocorrelation *τ*_ac_ of TC and RE population for spiking network (grey lines) and mean-field (blue/red lines).

**Fig. 3.**
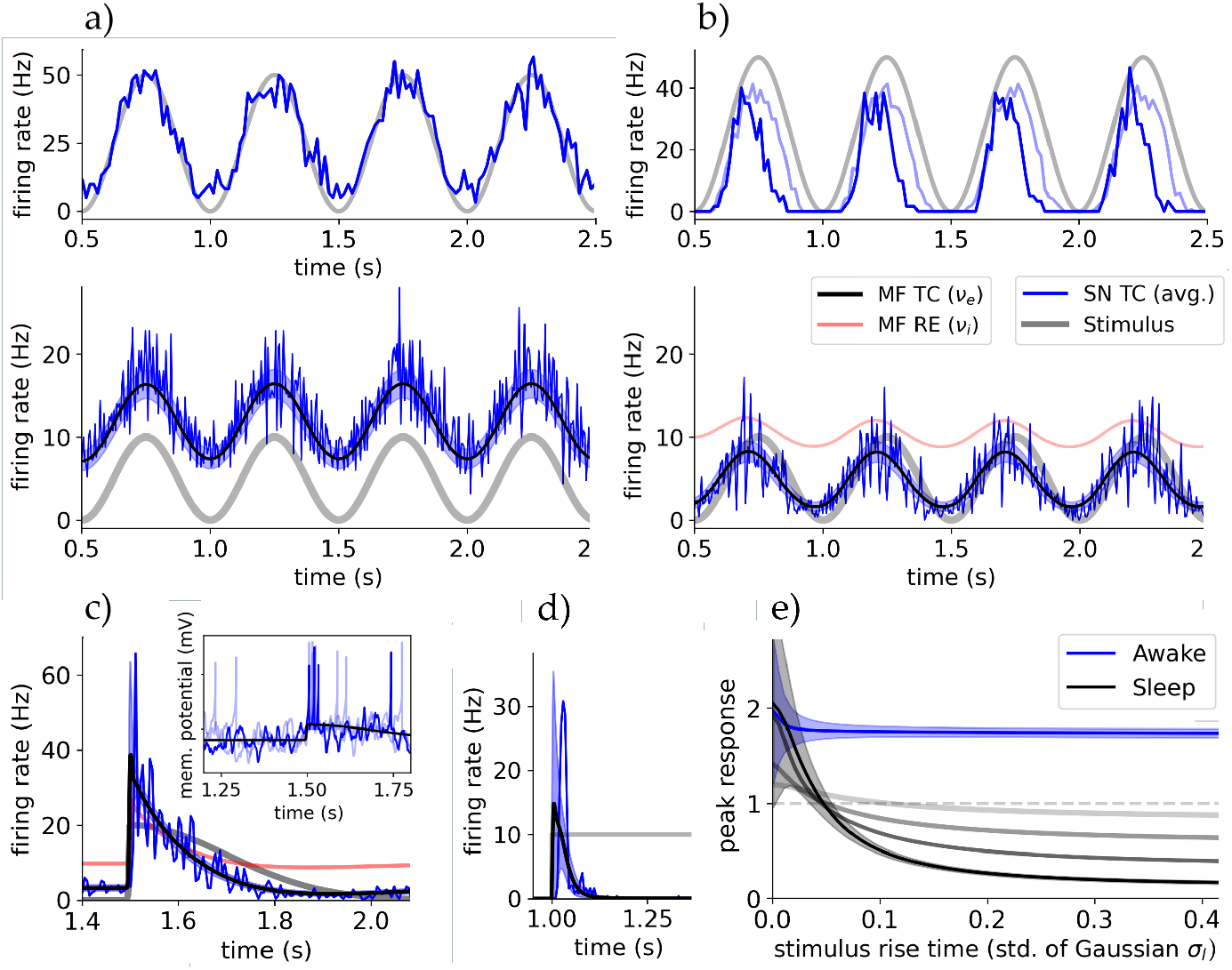
Bursting of TC cells renders thalamic response nonlinear in sleep state. **(a)** Top row: Single cell and population response to a strong oscillatory sensory drive *S* in awake state. Bottom row: Activities of spiking network (TC population, blue) and mean-field (TC population, black line with color-shaded std., RE population, red line). The grading stimulus is pictured in light grey. The single cell recording of the top rows is taken from this network simulation. RE activity is of same frequency and phase as TC activity but with amplitude ∼50Hz (not shown). **(b)** The same setup as in (a) but in sleep state (TC bursting, see main text). The single cell recording is done in the network of (a) which was in awake state to be close to the experiment. (Dark blue is sleep state and light blue is sleep state with lower adaptation *b* = 20pA.) The single cell traces in (a) and (b) reproduce the experiments of Sherman and Guillery [7, ch. 6]. **(c)** Thalamic response of spiking network and meanfield to a fast changing stimulus (split-Gaussian with steep left-hand std. *σ*_*l*_). Inset shows trace of 3 random TC cells of the spiking simulation, showing bursting at the onset of the stimulus (*t*_0_ = 1.5s; mV per s). **(d)** Rectangular stimulus in absence of cortical drive showing that TC activity vanishes after initial burst. **(e)** The maximum amplitude of response (peak), relative to the incoming stimulus amplitude, as a function of the ‘slope’ of stimulus (left std. *σ*_*l*_ of split-Gaussian; amplitude 10Hz and right std. 0.2s). Depicted is the response for awake and sleep state and in sleep state for different applied cortical constant drives (from black to gray: {1, 2, 4, 10}Hz).

A direct validation of the mean-field model is to compare the predicted mean firing rates and their standard deviations with those of the full spiking network, both modeling the entire thalamic substructure (Fig. 1a). This comparison is shown in Fig. 2c. Both populations receive an external constant *cortical* input of *P* = 4Hz and a split-Gaussian *sensory* stimulus *S* (definition in Section S.2). The spiking network provides the membrane potential evolution and spiking times *t*_*s*_ for all cells. The spikes of all neurons are shown in the upper raster plot. By averaging the number of spikes over a specific bin time *T*_bin_, we calculate the time-dependent averaged firing rate of the spiking population. We use *T*_bin_ = 5ms for all simulations except stated otherwise. To compare to the mean-field, in the formalism we have to employ a similar time window for the mean-fields time constant *T*, and we set *T* = *T*_bin_ (in accordance with the formalism requirements, Table S1). The spiking network and mean-field show the wanted balanced excitation-inhibition (E-I) state in AI regime with RE activity being dominant.

In Fig. 2d, we vary the cortical drive *P* and compare the equilibrium or stationary population firing rates (methodology in Section S.3) for TC and RE populations in both the spiking network and mean field over a 10-second simulation. This analysis reveals four distinct regimes of TC response: The first regime with no activity. The second regime with a fast response to changes in *P*. The *inhibited* third regime with limited responses. And the fourth regime with strong TC cell responsiveness due to (biologically unrealistic) saturated RE cell activity. This justifies using a cortical drive 1 *<P <* 10Hz for most simulations, ensuring a stable low-activity AI state, comparable to *in-vivo* experiments.

In Fig. 2e, we compare the distribution of firing rates and membrane potentials. In the latter the refractory states are removed to get a realistic comparison with the mean-field. The fit between mean-field and spiking network distributions only diverges at high firing rates of close to 100Hz due to the discontinuous nature of spiking models. The good agreement in not only firing rate but also membrane potential is significant, because equations (13) and (14) predict accurately the spiking populations membrane potential statistics and can henceforth be used to compare with electrophysiological data and methods.

In the same figure, top-right, there is depicted a comparison of (normalised) autocorrelations *τ*_ac_ of TC and RE population activity in the stationary state corresponding to *P* = 4Hz, showing a strong independence of population activity as expected from a inhibition-controlled network without excitatory-excitatory connections. This also agrees with the models being in AI state and the choice of *T* = *T*_bin_ = 5ms *> τ*_ac_ is justified.

Finally, we assess the robustness of the mean-field by varying global parameters (in Fig. S3). This is done for adaptation parameters {*b, a*}, which exhibit the significant change between awake and sleep state (Table 1), and synaptic excitatory conductance *Q*_*e*_ to validate its change for simulations in this study (Section 3.4). We demonstrate that even far of the actual fitting point, the mean-field remains effective in capturing network dynamics. This validation allows us to use the mean-field approach for parameter space analysis and the study of the transition between awake and sleep states with just one mean-field parameter fit.

### 3.2 Tonic and burst firing modes

We explore how bursting (the state of ACh neuromodulation) impacts the response of thalamic neurons and their network. Based on the fit to biological bursting TC cells from [44] we can already state that the employed parameter set with the AdEx shows *bursting* of single TC cells in the ACh-depleted or sleep state (as evident from the cell traces in Fig. 1).

We want to investigate the stability of those regimes and their dependence on model parameters. With the employed models (Section 2), the mechanism generating bursting is the slow adaptation current of the AdEx (1). In Section S.5 we derive an analytic metric quantifying firing adaptation using the transfer function of our meanfield framework. With this metric and with single-cell scans, we show in Fig. S1 that the awake and sleep states are well separated. While they are stable to small perturbations, they are also close to the phase transition which ensures richer dynamics.

Following, we aim to replicate experiments at single cell level on tonic and bursting states of TC cells, as documented in Sherman and Guillery [7, ch. 6]. These experiments involved manipulating the membrane potential of recorded TC cells to force either a tonic mode (around −65mV, resting state) or a bursting mode (around −75mV, hyperpolarised state), in the absence of external stimuli. A grating retinal stimulus was applied, leading to an oscillatory firing rate response. There, TC cells in tonic mode exhibited a linear response, while TC cells in bursting mode showed responses primarily during the initial phase of each stimulus period.

We recreated this behaviour computationally in our proposed spiking network with an oscillatory sensory drive *S*, with amplitude of 10Hz and frequency of 2Hz. The network was set in awake state emulating a lightly anaesthetised state as in experiment. To model the thalamus *in-vivo*, a constant external cortical drive of *P* = 4Hz was applied (the *inhibited* regime, Fig. 2d). Subsequently, we recorded one single cell with each awake and sleep parameters. While the proposed awake and sleep states are not identical to the artificially set tonic and bursting modes in the experiment, the switch via acetylcholine (ACh) generates a similar polarization.

The recorded cell’s response was calculated by averaging the spike times over 40 simulations for a time bin of 15s. This *firing rate* is depicted in the first row of each Fig. 3a,b for the awake state and sleep state, respectively. We observe the same response patterns as in the experiment for awake and sleep parameters, although with slightly lower response amplitudes in the sleep state compared to the hyperpolarized state of the experiment. This can be attributed to the absence of T-channels and low-threshold spikes in the AdEx model [7]. In Fig. 3b there is also depicted, in light blue, the response in sleep state with adaptation parameter in line with stated constraints (*b* = 20pA), which does not show the correct behavior.

Moving to population-level, the second row of Fig. 3a,b superimposes the spiking network’s and mean-field’s responses. In the awake state the entire TC population faithfully tracks the stimulus, as do single TC cells (top row). In sleep state, the response amplitude and also RE activity are greatly reduced, both showing phase locking while keeping the shape of the stimulus. The phase shift is created by the delay of slowly activating adaptation mechanisms and reactive RE inhibition. Phase locking was found in sleep state for all amplitudes of stimuli.

The effects, however, are quite small and functionally not so different between awake and sleep states. We would expect stronger effects of bursting in the responsiveness when adaptation effects are significantly slower than changes in the input and subsequently membrane potential (as is the case for single cells, Fig. 1b for a rectangular pulse). To investigate this at the network level, thalamic response to faster changing stimuli is tested. In Fig. 3c, a split-Gaussian with steep left-hand std. is depicted (at *t*_0_ = 1.5s with std. *σ*_*l*_ = 2*ms* and amplitude *A* = 20Hz, see Section S.1; inhibited regime). TC response is two-fold at an initial peak and then quickly adapts. This response curve is nonlinear and does not follow the shape of the stimulus faithfully anymore. This *peak* response is a direct effect of TC cells bursting at the onset of the stimulus, as shown in the inset for a random TC cell of the spiking network simulation. Similar to the single cells definition of showing bursting (Fig. 1b), also the TC population activity vanishes after the initial peak for a sustained input (no cortical drive, Fig. 3d). The initial bursting of TC cells is captured by the mean-field mainly via its second order moments, namely autocovariance *c* (blue shaded areas in plot) and autocorrelation *C* next to a smaller increase in mean firing rate *ν*.

To analyze the dependence of thalamic response for both tonic and bursting TC cells (awake and sleep state) on the shape of the stimulus, in Fig. 3e, there is depicted the *peak* response amplitude of the thalamus as a function of the std. *σ*_*l*_ of a split-Gaussian stimulus (*σ*_*r*_ = 0.2s and *A* = 10Hz), representing the change or ‘shape’ of a generic stimuli. In awake state the peak response is nearly constant, does not depend on how fast the stimulus changes, and the thalamus magnifies the input amplitude nearly two-fold. In contrast, in sleep state, only steep slopes or fast changing stimuli are generating a substantial response, whereas for slowly changing stimuli the response is drastically reduced (Fig. 3b).

In conclusion, both single-cell and population-level response of TC cells appears linear in awake state (ACh present) with enhanced stimulus amplitude, while in sleep state (ACh absent) response is linear but of reduced amplitude for slowly changing stimuli, and nonlinear for quickly changing stimuli. In addition, and as evident from Fig. 3c, both spiking network and mean-field model capture the bursting of TC cells, resulting in a ”bursting” population response. This enhances stimulus detection in low attention states for significant sensory inputs and the transmission of mostly time-dependent information such as oscillations in sleep state.

### 3.3 Cortical and sensory input

We will proceed with how thalamic responsiveness depends on background activity and how the two different biological inputs to the thalamus modulate it’s behaviour. Referring back to Fig. 3d, we see a modulating role of cortical input, which in sleep state can render the usually highly nonlinear TC response linear by removing the dependence on stimulus change at high cortical drives (gray lines in plot). This could allow the cortex to generate a time window where outside information temporally is transferred faithfully during usually non-attentive states.

Moving on, we are interested in the differences between the two drives. In Fig. 4a, the (stationary) firing rate response of the TC cell population in the mean-field model for different constant inputs in awake state is displayed, with both cortical and sensory drives. We applied a small constant cortical input *P* = 1Hz to be in a low activity AI state comparable to *in-vivo* (Fig. 2d). In case of sensory stimuli, the response is strongly proportional to the input, and we identify that the slope of this response is influenced by the cortical drive *P*.

**Fig. 4.**
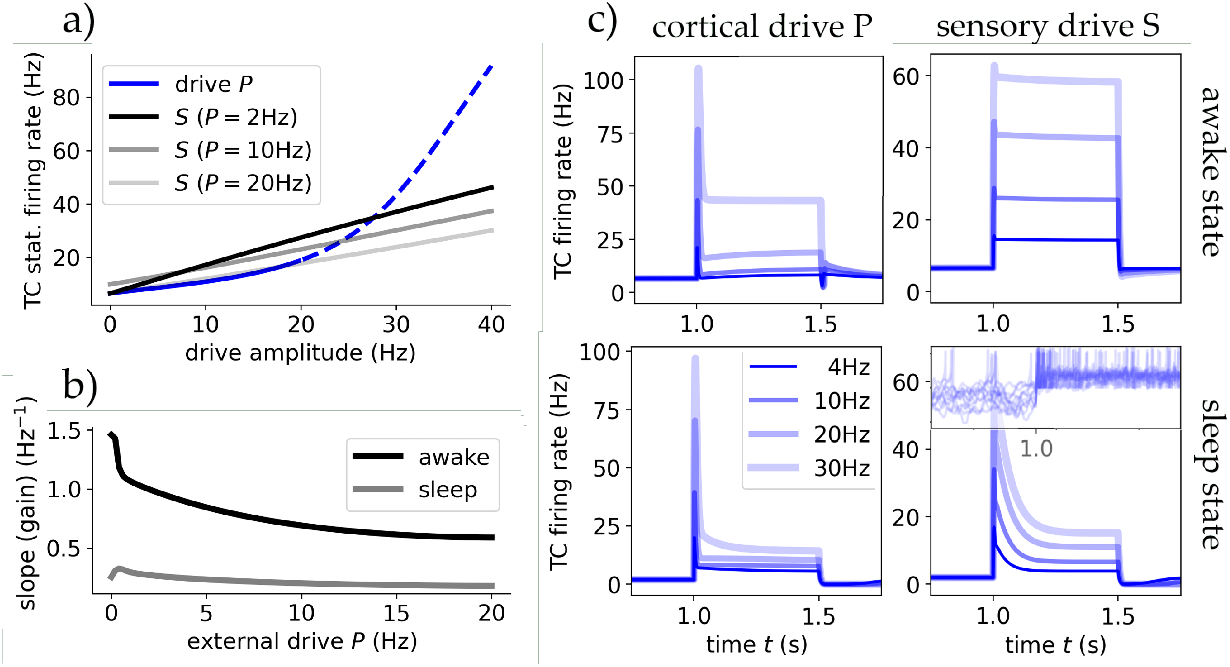
The thalamus’ responsiveness depends on external input origin. **(a)** The steady state output firing rate of the TC population after reaching equilibrium for different drives P and S. Blue is for inputs coming from cortical drive *P*, where solid marks the *inhibited regime* and dashed the *blow-up* regime. Black are for sensory drive *S*, with varying degrees of cortical input. TC cells respond linearly for sensory stimuli, whereas cortical stimuli are nonlinear only showing a strong proportionality after ∼20Hz. **(b)** Cortical drive removes thalamic response dependency on stimulus frequency. The gradients of the sensory input response curves (black in (a)) as a function of cortical input *P* for awake and sleep state. **(c)** The TC populations response to a rectangular stimulus of varying amplitude coming from either drive in both states. In the bottom-right there are also depicted 10 randomly chosen single cell traces to connect the population spike with single cell burst-like behaviour (mV per s).

In Fig. 4b we see this dependency is inversely proportional, where we conducted simulations for varying cortical drive amplitudes and observed that the gain (slope of the linear response curve) decreases as *P* increases. In the sleep state, the response remains relatively constant, slightly decreasing with *P*, contrasting the awake state’s high gain for all cortical inputs. The cortical drive removes the firing rate-dependency of thalamic response to stimuli in awake like states but does not alter it in sleep state. This is in agreement with studies which assumed the cortical role in the thalamus to be modulating thalamic response similar to noise [52], and with our study on synaptic noise (see next Section 3.4).

For cortical input, the response is nonlinear but exhibits multiple linear regions, as seen in Fig. 2c. The threshold at around 25Hz serves as a turning point (the end of the *inhibited* regime, at which the RE population firing rate saturates). Inputs below this threshold do not provoke a strong sustained response, while inputs above do. The RE population’s strong response to changes in the *inhibited* regime nearly nullifies TC and therefore thalamic response.

These behaviors are evident in the TC population’s response to a rectangular pulse stimulus from either *P* or *S* in Fig. 4c. Notably, low cortical inputs can even be repressive, with only larger amplitudes triggering robust and sustained responses, particularly in the awake state (in agreement with studies such like Crandall et al. [6]). In sleep state for both inputs or with low cortical inputs in awake state, responses are highly nonlinear, emphasizing the transfer of gradients rather than absolute values. The initial activity spikes at the onset of the input are created by the delay it takes the RE population to react to both stimulus and TC excitation to inhibit TC activity and –to a lesser extent– by the delayed adaptation mechanisms of both RE and TC populations. This is magnified in sleep state by stronger adaptation effects and resulting single cell bursting (see last Section 3.2). This mechanism allows the thalamus to respond to cortical input and modulation despite its strong inhibiting effect via the TRN.

Concluding, only in awake state and for sensory input, or with cortical control for sensory input at sleep state, thalamic responsiveness is linear while only temporal information is transferred for cortical input and sensory input at sleep states without cortical control.

### 3.4 Synaptic noise

We have analyzed so far how the responsiveness of the thalamic cells depends on the different firing modes and input sources. However, it has been shown that the level of synaptic noise (background activity) can significantly change these responses. We analyse in this section the role of noise as background synaptic and subthreshold activity and how it influences response and firing modes. We start by replicating single cell findings from Wolfart et al. [53]. They observed that synaptic noise controls TC neurons response and behaviour and that such noise removes the dependency of TC cells response on voltage and input frequency.

We recreated this computationally at single cell level. Fig. 5a shows the response of single TC cells in awake state to a Poissonian spike train of 5Hz with varying excitatory synaptic strength (*Q*_*e*_), reflecting the experimental setup. We observe the same step-like function in the static case without external synaptic noise: going from no activity to single spike response to double spike response or bursts at high conductances (regions separated by dashed lines). With noise the response function becomes smoother and the partition of the aforementioned regimes becomes blurred. The time-dependent noise was implemented as an Ornstein-Uhlenbeck (OU) process entering the cells membrane potential as a synaptic current (see Section S.2).

**Fig. 5.**
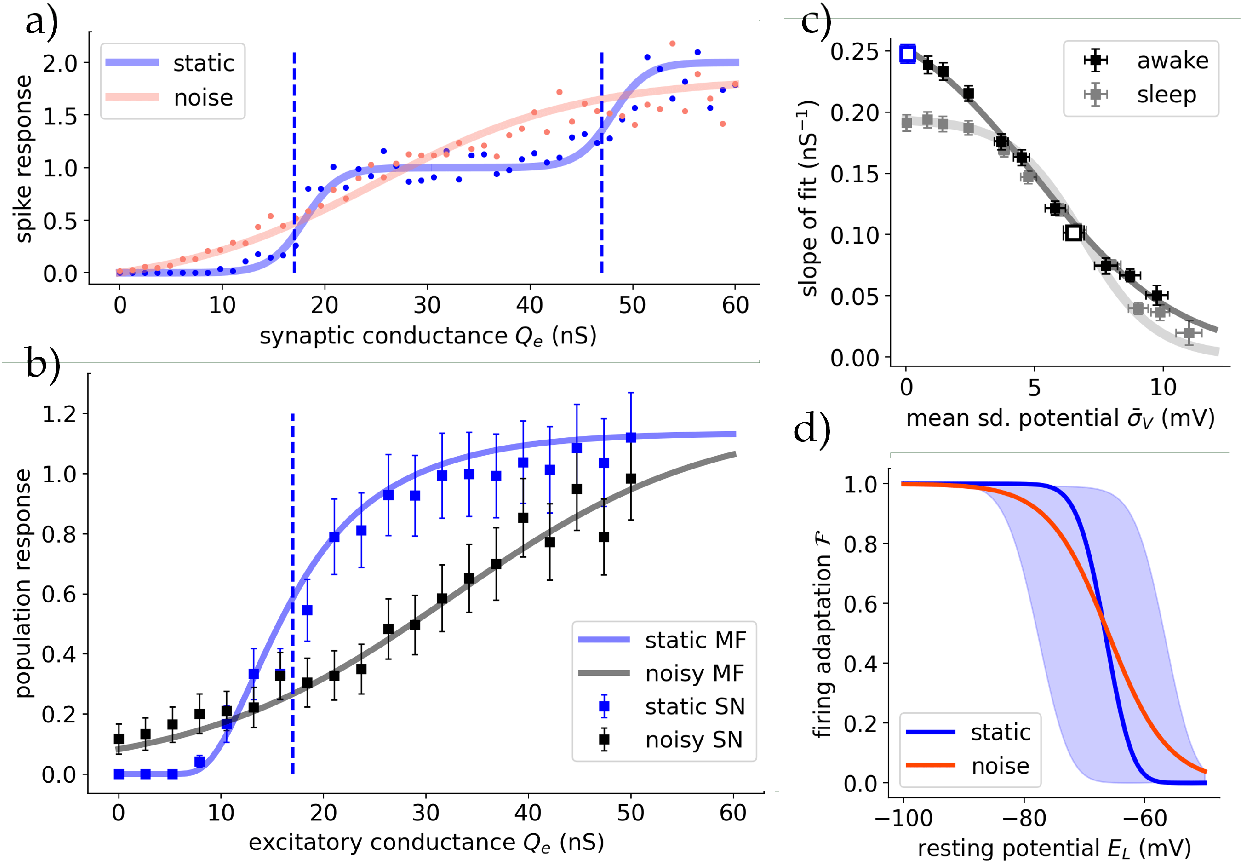
Synaptic noise modulates the dependence of thalamic responsiveness on voltage. (a) Response to a 5Hz Poissonian spike train for different values of excitatory synaptic strength *Q*_*e*_ for simulated spiking single cells in awake state. The noise was injected as an OU-like current in the membrane potential via *I*_syn_. The dots represent each the spikes per receiving incoming spike, averaged over 100 runs for 10s each. The lines correspond to sigmoidial fits, where the blue dashed lines mark the shift fit parameter depicting the center of the slope. Reproducing Fig. 5a of [53]. **(b)** The same setup but with the full thalamic spiking network (squares) and the mean field (lines), showing the relative response to a 10Hz Poissonian spike train. The dotted line marks the slope center of the single cell simulations going from no spikes to a one-to-one spike response. For the mean-field the synaptic noise was added as an additional time-dependent conductance into the formalism (see main text). **(c)** The (maximum) slopes of the mean-field response curves (b) plotted against the standard deviation of the membrane potential predicted by the mean-field. Showing a proportionality between fluctuations and the slope of the response function. At high noise levels the difference between awake and sleep state vanishes. The two cases from (b) are drawn as empty blue/black boxes. **(d)** Synaptic noise reduces the dependency of TC firing adaptation (sim. burstiness) on cell state (polarisation). Shaded area is the standard deviation induced by small conductance noise (5nS), and orange the average.

To translate this behaviour to the population level we did simulations of the full spiking network of the employed thalamic substructure. A constant Poissonian input of 15Hz was inserted into all cells, coming from just one source; comparable to dynamical patch clamps at single cell level. The stationary firing rate output of the TC population was measured for different synaptic strengths *Q*_*e*_. The resulting response function is depicted in Fig. 5b for the static and noisy case for both spiking network and mean-field.

For the mean-field, the noise-dependent shape of the response function is passively included in the definition of the transfer function (8), with its slope being controlled by the standard deviation of the subthreshold membrane potential (*σ*_*V*_). However, to recreate the experiment, which employed a time-dependent external noise, we extended the formalism by adding two additional static synaptic conductances 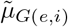. Those are modelled as OU-type functions averaged for each time bin equal to the mean-field’s time constant *T* (see Section S.2).

Both spiking network and mean-field show that the TC populations response function has its maximum slope at the same place as the first step at single cell level from no activity to single spike response (the first dashed line in Fig. 5a and the dashed line in Fig. 5b, respectively). Furthermore, the effect of synaptic noise is the same for population response as in the single-cell experiment, decreasing the response functions maximum slope (see Fig. 1c in [53]).

How the maximum slope of the response function depends on this noise is depicted in Fig. 5c (compare to single-cell experiment; Fig. 1c inset in [53]). Here instead of the injected noise the noise-dependent membrane potential subthreshold fluctuations averaged over all runs 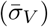 is shown. In sleep state the population response slope is ∼20% less steep for the static case or small noise. Strong synaptic noise and subsequent membrane potential fluctuations decrease the slope as expected. Additionally, synaptic noise diffuses the response differences of awake and sleep state at intermediate noise levels and removes nearly all dependence of thalamic response on conductance at high noise levels 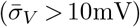, where the response function is nearly constant (at a value dependent on the ratio of excitatory and inhibitory noise 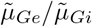.

Additionally, the effect of synaptic noise on the firing adaptation *ℱ* of TC cells was tested in Fig. 5d. Noise diffuses the state transition between no firing adaptation and strong firing adaptation for different levels of membrane potential polarization. As in Section 3.2 we can refer to the similarity of *ℱ* to *burstiness*, and hypothesize that strong noise allows for firing adaptation and also bursting for membrane potential levels showing no bursting without noise. Although the effect for the noise studied here is smaller, this is qualitatively in agreement with experimental study ([53] Fig. 5b therein).

Previously, we showed that synaptic noise modifies thalamic response dependency on voltage and conductance. There, input frequency was fixed. Further following [53], we proceed to investigate how noise changes the thalamus’ response in respect to input frequency.

For this, single TC cells were simulated for extended duration with incoming Poissonian spike trains of 10Hz, modelling a generic input from retinal ganglion cells in-vivo. Here the *retinal* input conductances were fixed. During simulation, for each output spike of the recorded TC cell, the interspike interval (Δ ISI) of the retinal input between the spike which results in the spike response and the preceding one is measured. This way the spike probability or response can be measured as a function of input frequency. Because of the increasingly more rare occurrence of large ISI’s (Δ *>* 400ms) in a Poissonian spike train of 10Hz, the following plots are cut of at 550ms. Until then reasonable long simulation times provide distinguishable uncertainties. Fig. 6a shows the results for a TC cell without additional synaptic noise. At resting potential (awake state, *E*_*L*_ = −65mV with *Q*_*e*_ = 14ns) spike response only occurred at summed input spikes with Δ *<* 50ms with an all-or-none character. At hyperpolarized potential (awake state, *E*_*L*_ = −70mV with *Q*_*e*_ = 24ns) not only input spike summation evoked a response but also ISI’s with duration longer than 300ms. These even show higher spike probability compared to spike summation at low ISI’s. The difference in input conductances *Q*_*e*_ was necessary to account for equal number of spikes between both states, where the high conductance in the hyperpolarized state captures the effects of T-channels. In the presence of synaptic noise this changes drastically and both TC cells at resting and at hyperpolarized levels exhibit the same spike response, completely independent of input frequency (see Fig. 6b). Remarkably, with noise spike probability is significantly lower even with spike summation (ISI → 0ms), independent of polarization. These results correctly reproduce the experimental results of [53].

**Fig. 6.**
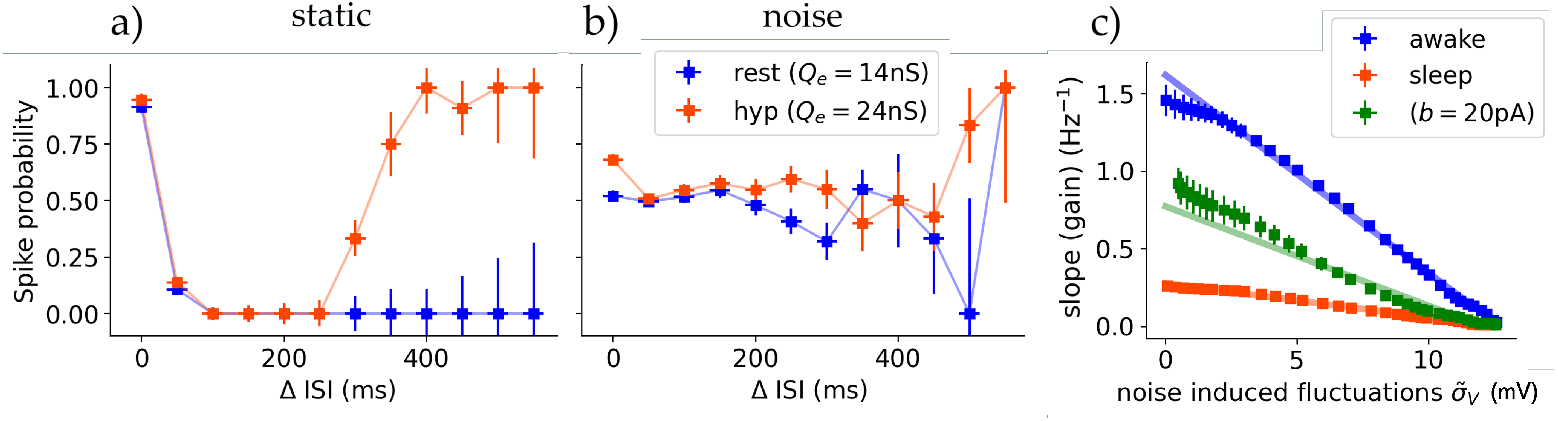
Synaptic noise removes the dependence of thalamic response on frequency. **(a)** A single TC cell’s spike response probability dependent on the interspike intervall (Δ ISI) between input spikes. The input is a Poissonian spike train with a mean frequency of 10Hz, comparable to an in-vivo retinal input. For both resting (*E*_*L*_ = −65mV, blue curve) and hyperpolarized (*E*_*L*_ = −70mV, orange curve) states the spike response is nearly 100% at low ISI’s and therefore only reacting to *summed* input spikes. In hyperpolarized state, with T-channel adjusted synaptic conductance (see main text), the TC cell responds to also high ISI’s with a nearly one-to-one spike probability. This reproduces Fig. 4 in [53]. **(b)** Same setup as in (a) but with additional synaptic noise (see main text). Frequency-dependent response is nearly removed. **(c)** Thalamic stationary response slope (gain per increase of input, see main text) of the mean-field to gated sensory stimuli of 10Hz as a function of synaptic noise via noise induced membrane potential fluctuations. For the awake state, and the sleep state with *b* ∈ {20, 200} pA. Regardless of state, noise linearly leads to a reduced thalamic response dependency on stimulus frequency. The shaded lines are linear fits.

Moving to population level, we present thalamic stimulus response as a function dependent on synaptic noise. Noise acts in a similar way on the frequency dependent response as cortical input (see Section 3.3). In the same manner, in Fig. 6c the slope of the linear response of the TC population as a function of input amplitude is depicted (gain). As with modulating cortical input (refer Fig. 4), noise decreases response.

However, different to the control of cortical input, where the gain saturates at 0.7Hz^−1^, noise linearly reduces gain until a complete banishment of frequency dependence at very high noise levels 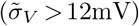. This holds true for all states. In the awake state the loss of gain per membrane fluctuation is (−0.12 ± 0.01)gain/mV. For the sleep state the loss is (−0.028±0.003)gain/mV. Finally, we see that the noise required to equalize the dependence on frequency between awake and sleep state is significantly higher than for equalizing the dependence on voltage (induced subthreshold fluctuations of 12mV and 4mV, respectively).

### 3.5 Spindle oscillations

Spindle oscillations are one of the main activity dynamics of the thalamus during NREM sleep or anesthesia [54], strongly influencing the responsiveness of the thalamus in such states. These originate from the superposition of multiple cellular and circuit properties, with especially the mechanism of RE-induced rebound bursts in TC cells in ACh depleted or sleep-like states (see Section S.6 for more details).

To enhance this rebound bursting in our sleep state we promote burst firing by adjusting the reset membrane potential (*V*_*r*_) below the sodium spike threshold onset: *V*_*r*_ = −48mV for TC and *V*_*r*_ = −42mV for RE cell (see [29] for the significant role of *V*_*r*_). This yields sustained burst firing without sustained activation, mimicking T-channel like activation and IPSP barrages in RE cells, which we could not capture with the AdEx sleep state (Section 3.2). Accordingly we re-calibrate the mean-field fit to accommodate the change in *V*_*r*_ (Table S2).

We observe spindles in the proposed models within this adjusted sleep state and when applying an initial kick to evoke activity. Fig. 7a,b show self sustained oscillations of both full spiking network and mean-field, respectively. Their frequency spectrum and phase space are compared in Fig. 7c.

**Fig. 7.**
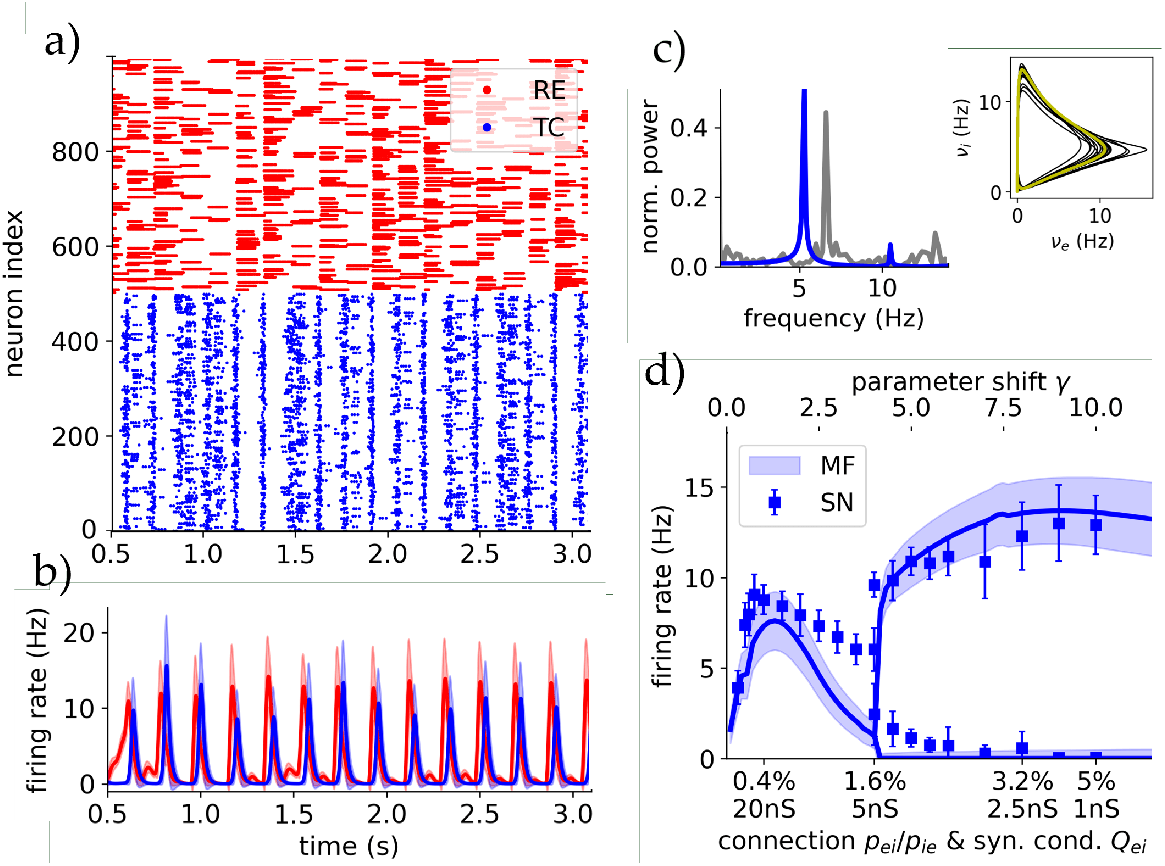
Spindle oscillations in a sleep-like state generate a highly unresponsive thalamic state. **(a)** Raster plot of the full-scale spiking network with 1000 neurons for spindle parameters (ACh/sleep state with rebound burst). **(b)** Mean-field oscillations: Firing rates and standard deviations of both TC and RE populations. **(c)** Fourier spectrum for spiking network (grey) and mean field (blue) of the TC population activity. Inset right: Phase plane in the TC and RE firing rate space. Yellow is the stable limit cycle and black the transient. **(d)** Bifurcation diagram showing the suggested Andronov-Hopf bifurcation that occurs when gradually increasing the connection probability in the network, for spiking network and mean-field. This corresponds to a parameter shift from the parameters used in [31] with *γ* = 1 to the parameters used in this paper with *γ* = 10.

In a previous study [31] only small AdEx networks generated spindles in SR-like dynamics, while at larger scales of *N >* 40 neurons, population activity showed self-driven steady states with AI dynamics. In Fig. 7d we show a bifurcation diagram transitioning between connection and balancing synaptic parameters from [31] (*γ* = 1) to our parameter values (Table 1, *γ* = 10). Increasing connection probability creates a supercritical Andronov-Hopf bifurcation, showing that sufficient connections are necessary for generating and keeping stable spindles at larger network scales. The spindles produced by our mesoscale network show realistic SI dynamics (see Section S.6).

This self-sustained oscillation is remarkably robust in regards to perturbations of all kinds of inputs, producing spindles of same frequency. This renders the thalamus’ responsiveness in this spindle-adjusted sleep state highly independent of external input. Only prolonged and constant inputs of a duration longer than multiple spindle periods destroy the synchronisation and create steady state AI dynamics. However, spindle oscillations emerge again as soon as the input stops. The bifurcation diagram in connection with [31], shows that thalamic function and responsiveness can be drastically altered depending on specific order parameters, as seen here with connection probability and synaptic conductance.

## 4 Discussion

In this study, we investigated the state-dependent responsiveness of the thalamus at micro to meso scale. For this we introduced a biologically realistic mean-field model of the thalamus, which captures the population dynamics of thalamocortical relay neurons (TC) and thalamic reticular neurons (RE) in two physiological states: Awake state (high level of ACh neuromodulation, wakefulness and REM sleep) and sleep state (low level of ACh neuromodulation, NREM sleep; Section 2.2 and [9, 44]).

The mean-field model employs the master-equation formalism introduced by El Boustani and Destexhe [48] and incorporates adaptation mechanisms [22]. We constructed it using a bottom-up approach following the formalism described by Zerlaut et al. [51], which includes a subthreshold-dependent transfer function [50].

In Section 3.1 we successfully validated the mean-field’s predictive accuracy through comparison with the spiking network, confirming its ability to replicate the dynamic behavior and population distribution of thalamic cells. We also demonstrated that the mean-field model is capable of predicting the network’s subthreshold activity and proved its validity beyond the fitting point. This allows the use of modeling experiments using intracellularly injected currents in combination with this model.

Thalamic responsiveness and it’s dependence on internal and external state was investigated in three steps: First, in Section 3.2, we analyzed the important role of bursting in TC cells which provides a mechanism by which the thalamus modulates the transmission of sensory information to the cortex, extending the single cell findings of Sherman and Guillery [7]. We showed that in sleep state response is highly reduced, except for significant (fast changing) stimuli where mainly their timing is transmitted via a strong and fast thalamic response, which is generated by TC cell bursting and delayed inhibition of RE cells. This is in agreement with the hypothesis that bursting plays a role in generating *wake-up* calls during low-attention states [7], and also supports that the thalamus generates and distributes oscillations in NREM sleep states [54]. Additionally, an important validation of the proposed mean-field model as a *thalamic* model is that it captures bursting dynamics, a defining thalamic feature. This is also nontrivial as bursting is highly nonlinear and spike-time dependent, whereas the model is firing rate-based, suggesting interesting future theoretical work.

Second, in Section 3.3, we examined the influence of external states on thalamic response. We demonstrated that in this model, and in accordance with experiments [6], there is an important distinction in the origin of inputs: sensory-like stimuli experience a more linear response and are therefore transferred more faithfully than cortical-like inputs, which generate a nonlinear response. In sleep-like state the relay of information becomes strongly nonlinear regardless of input origin. Additionally, we identified the modulatory effect of cortical input to (1) repress thalamic response in awake state, via activation of the inhibiting TRN, and (2) to promote a linear response to sensory stimulus in sleep state. (2) would allow the cortex to *wake-up* the thalamus in order to faithfully transfer sensory input, e.g. after a preceding wake-up call of a potentially significant stimulus.

Third, in Section 3.4, we investigated the role of synaptic noise in thalamic response. The experimental findings of Wolfart et al. [53] were as a first time successfully modeled. We showed that synaptic noise acts as a controller for response also at the population level. The TC cells’ step-like response function for single spikes translates well into their collective response at population scale, sharing the same conductance threshold. This allows the thalamus to fine-tune its responsiveness to external stimuli at cell and population level. Additionally, noise diffuses transitions between states of tonic/bursting firing at single cell level and awake/sleep at the population level. We find that in equal manner for single cell and population level, noise banishes the thalamic response dependency on both voltage and frequency. We state the interesting similarity between synaptic noise and cortical input in how both control stimulus transfer and render stimulus response less dependent on stimulus frequency, whose similarity is often only presumed [52]. These insights pronounce the importance of integrating conductance-based subthreshold fluctuations dynamics into meso to macro scale modeling approaches.

Finally, in Section 3.5, the successful reproduction of spindle-like oscillations in a sleep-like state is an important validation for our thalamic model. We emphasize the necessity of specific substructures within the thalamus for generating realistic oscillations at all scales ([55], see Section S.6). In this state thalamic responsiveness to inputs is highly suppressed. Only strong and prolonged cortical inputs temporarily create AI dynamics during their activation.

In conclusion, our study underscores the value of integrating single-cell dynamics with thalamic specific structure at population-level in understanding the complex role of thalamic responsiveness. With these findings and with offering a biologically realistic and experimentally grounded mean-field model of the thalamus, which captures the effects of bursting, neuromodulation, and fluctuation, we provide here an essential starting point for: (1) Further investigation of thalamic function and sensory processing. (2) Large-scale modeling (especially the thalamo-cortical loop with already developed cortical mean-fields [22, 34]), while integrating micro-scale cell and synaptic effects with physiological states.

## Data Availability

The numerical codes for simulating mean-field and spiking network and the code for fitting the transfer function of the mean-field on single cell data are openly available at the corresponding `GitHub` page: https://github.com/joverwiening/Thalamic-meanfield.

## Author declarations

The authors declare no competing conflict of interests.

## Acknowledgments

Research supported by the CNRS and the European Union (Human Brain Project H2020-945539 to AD, Virtual Brain Twin project Horizon Health 101137289 to AD) and the ANR (ImpactCom project to AD). The funders had no role in this study.

## S Supplementary

### S.1 Additional formulae

#### Transfer function

The transfer function *F* was defined as the complementary error function

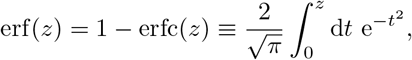

With erf : ℝ → (−1, 1). The statistical interpretation is that for a normal distributed random variable *ξ* with ⟨*ξ*⟩ = 0 and 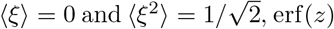, erf(*z*) is the probability of *ξ* ∈ [−*z, z*].

#### Mean-field correlation

The differential equation describing the correlation of populations {*µ, ν, λ*} ∈ {*e, i*} to close the second order statistical moments of the mean-field formalism is given by

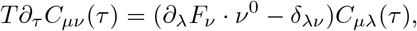

where *τ* is the time lag and *ν*^0^ marks the stationary solution of the firing rate.

### S.2 Details on stimuli

#### Gaussian stimulus

The *split-Gaussian* stimulus employed as a general sensory input was defined as

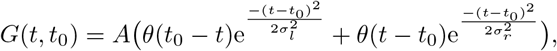

Where *θ*(*t*) is the Heaviside step-function, and with *A* the amplitude and *σ*_*l,r*_ of the slopes he left and right side of the split-Gaussian, respectively.

#### Synaptic noise model

Ornstein-Uhlenbeck type noise can be defined as

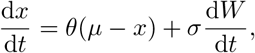

where *µ* is the mean reversion level, *θ* the mean reversion rate, and *σ* the amplitude of the standard Wiener process *W* (*t*), or white noise.

The conductance noise was implemented in the mean-field via adding the mean of OU-type noise (*x*) in a time window corresponding to *T* to the static conductances *μ*_G_ (see methods). The resulting conductances with including noise are then:

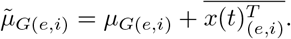

The specific parameters for *x*(*t*) used for the case of synaptic noise case in the simulations are *µ* = 0nS, *θ* = 5ms^−1^, and for the amplitudes *σ*_*i*_ = 200nS and *σ*_*e*_ = 60nS.

### S.3 Dynamical analysis

The steady and equilibrium points of the spiking network are taken as the averaged population activity over long simulation times for constant inputs where oscillations are ruled out either by observation or by spectrum analysis. For the mean-field these are calculated as the populations activity after a transient up to a point where the Jacobian (*J*) eigenvalues are negative. The Jacobian is defined on the variable set *X* ∈ {*ν*_*e*_, *ν*_*i*_, *ω*_*e*_, *ω*_*i*_} and calculated via numerical derivatives.

For the bifurcation diagram (Fig. 7) the same technique was used up to the bifurcation point, except that now 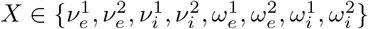. For the mean-field the Jacobian eigenvalues are complex up to the bifurcation point, suggesting a supercritical Andronov-Hopf bifurcation. After the bifurcation point for the spiking network, instead of the averaged firing rates, the extrema are averaged over a long time simulation, mimicking the real sub-plane of the suggested complex-space Andronov-Hopf bifurcation. The same was done for the mean-field.

The maximum Lyapunov exponents are calculated employing the Rosenstein algorithm [56].

### S.4 Numerical simulations

All simulations were conducted in `Python`. The spiking network simulations additionally used the `Python` package `Brian2` [57]. Constructing the network, all neuron connections to random to ensure a statistical model comparable to the mean-fields assumptions. All numerical integrations in spiking network simulations employed *Heun’s method*, while for mean-field simulations *Euler’s method* was used.

### S.5 Firing adaptation

We introduce firing adaptation *ℱ* as a metric describing how adaptation shapes the response. Because the AdEx generates bursting-like behaviour via its adaptation current *ω*, we will see a good agreement between *ℱ* and burstiness.

We use the transfer function *F* (8). For the effect of adaptation mechanisms on the firing rate of a neuron type we can identify two crucial states described by *F* : The no-adaptation fixpoint (*F*_0_): This state represents the firing rate of a cell in the absence of adaptation and corresponds to the cell’s firing rate at the onset of a stimulus where the (slow) adaptation mechanisms have no impact yet. And the real-adaptation fixpoint (*F*_*ω*_): Representing the cell’s firing rate when it has fully adapted to its own and the stimulus influence. Then *F*_0_ − *F*_*ω*_ reflects the change in firing rate of a cell from initial activity at the onset of a stimulus as it transitions towards full adaptation and slowing its firing rate.

Necessarily, the transfer function is firing-based and needs non-zero firing rate inputs (*ν*_*e*_, *ν*_*i*_) to yield results. Therefore, we calculate the fixpoints for a constant *ν*_*e*_ = 1Hz. When considering high firing rates as decreased ISIs this metric is comparable to experimental methods measuring burstiness (such as [53]). Concluding, we define the firing adaptation metric as

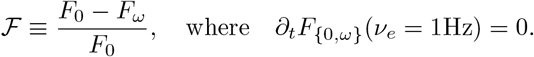

With this, we investigate the stability of bursting in TC cells and the dependence of bursting on model parameters. For this we use the level of firing adaptation to quantify the effect of adaptation mechanisms on the firing rate of neurons. The dependence of TC firing adaptation on membrane and spiking adaptation parameters and membrane polarization is shown in Fig. S1a,b. Firing adaptation is strongest at high membrane adaptation levels and hyperpolarized membrane potentials. The awake state experiences nearly no firing adaptation, while the sleep state shows strong firing adaptation.

**Fig. S1.**
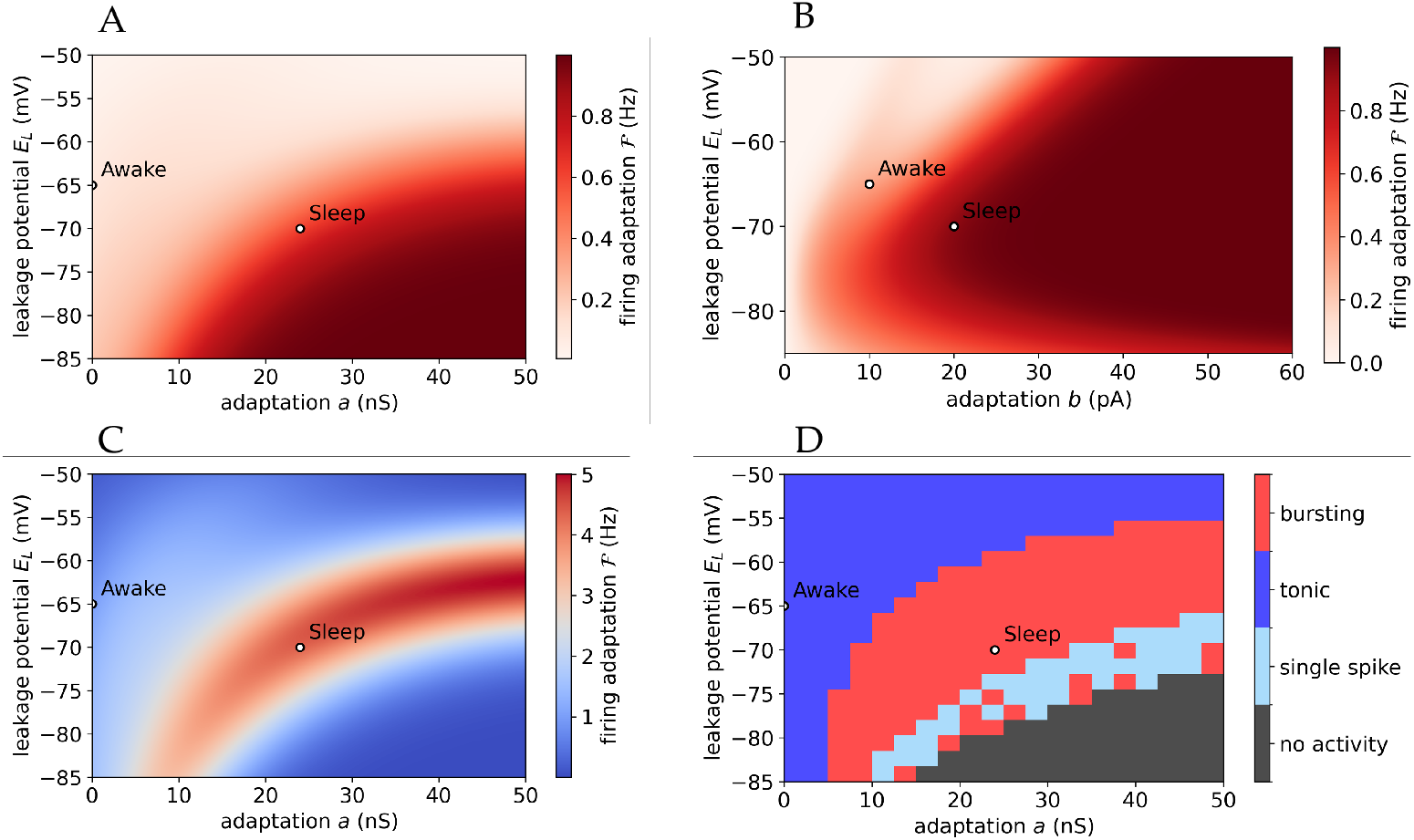
Firing adaptation and dynamics of TC cells. **(a)** Parameter scan for firing adaptation *ℱ* for TC cells via the fitted transfer function (8) for resting potential and membrane adaptation a. Depicted are the parameter values for sleep and awake state. **(b)** Same as in (a) but for spiking adaptation b. **(c)** The scan of (a) but the firing adaptation *ℱ* is multiplied by the firing rate during the response (also via transfer function), revealing a split of bursting dynamics for the same input. (d) Parameter scan of TC single-cell simulations, showing the possible firing modes of tonic, bursting, single spikes, or no spikes as a response to a input current (see text). Similarity to firing adaptation in (a)) and (c) is evident.

We want to improve on defining the states of ACh as tonic/bursting states and subsequently as awake and sleep states. For this we conducted a parameter scan using spiking network simulations of single TC cells mapping the different firing modes. Similar to [58] (and Fig. 1) we injected the cell with a constant current for 1s. We classified firing patterns based on the number of spikes on a time scale relative to the adaptation time constant (*τ*_*ω*_ ≃ 200 ms). The external current applied was proportional to *E*_*L*_ in order to induce activity (with *I* = {200nA for *E*_*L*_ = −50mV, 400nA for *E*_*L*_ = −85mV}). Fig. S1d presents the results, highlighting the four possible firing modes. The ACh-absent or sleep state exhibits stable bursting not susceptible to either adaptation or voltage perturbations while the ACh-present or awake state is deep in the tonic regime.

The scan shows a similar tendency of increased bursting as with increased firing adaptation. Note that the firing adaptation is not taking into account if there is actually a non-zero response to account for the single spikes or no activity response types of the single cell scan (Fig. S1d). When integrating with *ℱ* the actual response amplitude we get however the same strip-region of bursting as in the single-cell scan (see Fig. S1c, the same holds for the scan in *b*).

### S.6 Spindle mechanism

Spindle oscillations are one of the main activity dynamics of the thalamus during slow-wave sleep or anesthesia [54], strongly influencing the responsiveness of the thalamus in such states. These originate from the superposition of multiple cellular and circuit properties. In ACh depleted conditions such as during sleep, T-channel currents promote bursts after hyperpolarization (as modeled in [59] for a Hodgkin-Huxley model), and barrages of IPSPs occur in strongly connected subgroups within the TRN [60]. This creates two self-sustained loops: an inhibitory self-loop within TRN neurons, and loop connecting RE and TC cells (see colored connections in Fig. S2e). The rebound bursts caused by this circuit arrangement generate spindle oscillations, as shown by experiments and computational models [54, 61].

Computational models demonstrated that a 4-neuron structure of TC and RE neurons exhibits self-sustained oscillations with all characteristics of spindles, as shown first for Hodgkin-Huxley type models [62] and later using AdEx models [31]. These 1015 oscillations rely on a loop where a TC cell triggers a burst in an RE cell which, in turn, causes a rebound burst in another TC cell, completing the cycle. In their study however the oscillations waned with increased network size.

In this study, we showed that the proposed models in a sleep-like state indeed show spindles at all scales, validating them as thalamic models. In contrast to a previous study [31], we aimed for robust spindles at population scales. To enhance rebound bursting we promoted burst firing by adjusting the reset membrane potential (*V*_*r*_) below the sodium spike threshold onset: *V*_*r*_ = −48mV for TC and *V*_*r*_ = − 42mV for RE cell. This yields sustained burst firing without sustained activation, mimicking T-channel like activation and IPSP barrages in RE cells. Accordingly we re-calibrated the mean-field fit to accommodate the change in *V*_*r*_. (The original fit produces spindles with unrealistic amplitudes.) The resulting fit parameters are depicted in Table S2.

The Jacobian eigenvalues calculated at the steady points of the mean-field up to the bifurcation point are complex. This together with the transition into a stable cycle suggests a supercritical Andronov-Hopf bifurcation as the origin of spindles Fig. 7d).

**Fig. S2.**
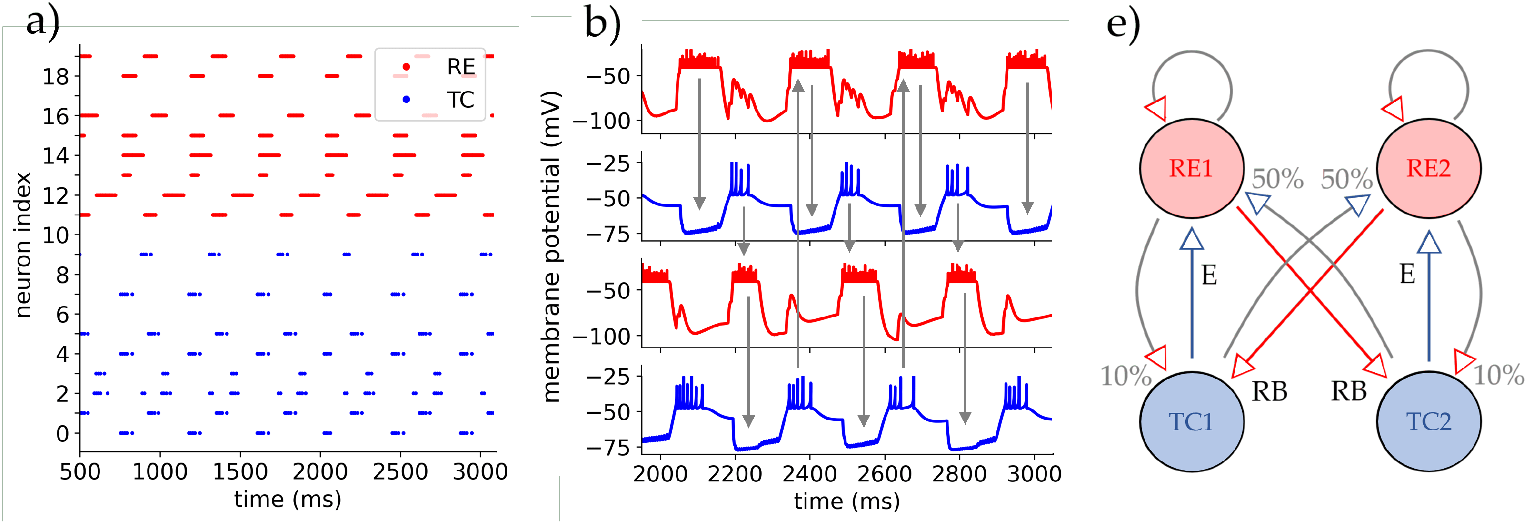
Spindle oscillations are generated by sparse TC-RE connections. **(a)** Raster plot of a simulation for the small network showing all 20 neurons. **(b)** Four neurons showing anti-phase burst firing with rebound bursts of the TC cells as a possible generator of spindles. Cell traces are taken from the simulation in (a) with a small network (*N* = 20). The arrows show the loop of activity propagation between RE and TC cells with rebound bursts and direct excitation. These pathways translate to the four colored connections in (e). **(e)** The 4-structure used for the mean-field connections. The colored arrows mark the main pathway of propagation with rebound bursts (RB) and direct excitation (E) as transmitters. The non-significant connections are marked in grey with their respective relative strength shown in percentages of the original connection probability. The RE populations keep their local self-loop.

In Fig. S2a oscillatory dynamics in a small network of 20 neurons are illustrated, revealing a generating 4-loop structure inside the network (b). We employed the burst-adjusted sleep parameters but with synaptic parameters from [31]. To invoke activity, an initial kick of Poissonian input of high frequency to only a random subset of neurons was applied.

For the mean-field to produce self-sustained oscillations we employ the 4-structure as seen at single cell level (from [31]) and justify its need also at the population level. This is further supported by studies such as [46]. They showed the mechanism behind the generation and specific frequency of oscillations in the thalamus is the delay in the propagation of activity between different TC neurons. Thalamic relay cell clusters have no connection to themselves and so RE neurons serve as intermitter. The slow adaptation mechanisms governing rebound bursts are therefore generating the slow ∼10Hz spindle oscillations.

In order to implement this, we construct two RE and two TC populations and keep all connections active, but reduce the connection probability of links which are antagonistic to this propagation via rebound bursts. This can be seen as two close but locally separated subgroups of relay neurons. The resulting structure is depicted in Fig. S2e. (Note: If the connections are set homogeneous then this network is acting exactly like a mean-field with just two populations.) The main links for the propagation are colored and labeled with their mechanism of propagation: RE induced rebound bursts and TC excitation. To initiate the spindles here also a kick of high activity going to only one or two populations was necessary, in agreement with the spiking network. Additionally, *T* was set to 15ms to have similar phases and shapes with the spiking network.

The spindle-adjusted sleep state differs quite substantially from the sleep state in the rest of this study. The adapted leakage and reset parameters, T-modelling channel like activation and IPSP barrages to invoke stronger rebound bursts, create this significant difference, suggesting two distinct states. One representing a sleep state similar to N2 in which spindles are observed, while the other could represent a less-deep sleep state [2, ch. 44]. This suggests that the concentration of ACh alone is not sufficient for investigating awake and sleep separation, but the mechanisms behind rebound bursts are specifically important to include. Especially how these connect to ACh and other, in this study neglected, neuromodulators like dopamine, norepinephrine, or serotonin. Additionally, we remark that the shown spindles inherit the correct underlying mechanisms, as evident from experiments, but do show a frequency at the lower end of the usually defined spindle frequency, potentially being more close to Delta waves. All this hints at interesting future work in connecting physiological brain states of awake and sleep with multi-scale thalamic models.

### S.7 Supplementary Tables and Figures

**Table S1.** Formal requirements and restrictions for the employed approximations of the mean-field formalism.

|  |  |
| --- | --- |
| $\tau_\omega > \tau_m$ | Mixed activity regime of the AdEx [29] |
| $T > \tau_{ac} \approx 1\text{ms}$ | No fluctuations (see Fig. 2c), with $\tau_{ac}$ the autocorrelation time |
| $T < \max(\nu_{AI})^{-1} \approx 15\text{ms}$ | with $\nu_{AI}$ the maximum firing rate in the thalamus [7] |
| $\Delta\nu = (NT)^{-1} \simeq 0.1\text{Hz}$ | mean-field resolution [48] |
| $\tau_\omega \gg T$ | $\omega$ dynamics are independent of fluctuations in $\nu$ [22] |
| $\tau_m > \tau_{(e,i)}$ | dynamics of $v(t)$ are stat. independent of synaptic changes |
| $\mu_G^s/\sigma_G^s \propto \sqrt{\nu_s} \ll 1$ | approximation for synaptic fluctuations (10)&(11) [50] |
| $\langle\tau_{\text{eff}}\rangle/\sigma_\tau \propto \sqrt{\sum_s \nu_s} \ll 1$ | approximation for membrane time constant [50] |
| Sparse and random network | $G(N, p)$ Erdos-Renyi model with $p \approx 1/N$ [48] |
| Activity regime | Asynchronous Irregular (AI) in E-I balance [49, 63] |

**Table S2.** The fitting parameter values for recreating spindles with sleep parameters and included rebound burst mechanism. (All values in mV.)

| cell | $P_0$ | $P_\mu$ | $P_\sigma$ | $P_\tau$ | $P_{\mu\mu}$ | $P_{\mu\sigma}$ | $P_{\mu\tau}$ | $P_{\sigma\sigma}$ | $P_{\sigma\tau}$ | $P_{\tau\tau}$ |
| --- | --- | --- | --- | --- | --- | --- | --- | --- | --- | --- |
| TC | -51.17 | 3.94 | 15.53 | -7.15 | 0.35 | -7.57 | -1.19 | -13.61 | 9.47 | 29.44 |
| RE | -45.84 | 3.53 | -16.90 | 41.75 | 0.34 | 2.02 | -5.23 | 19.44 | 49.70 | -93.53 |

**Fig. S3.**
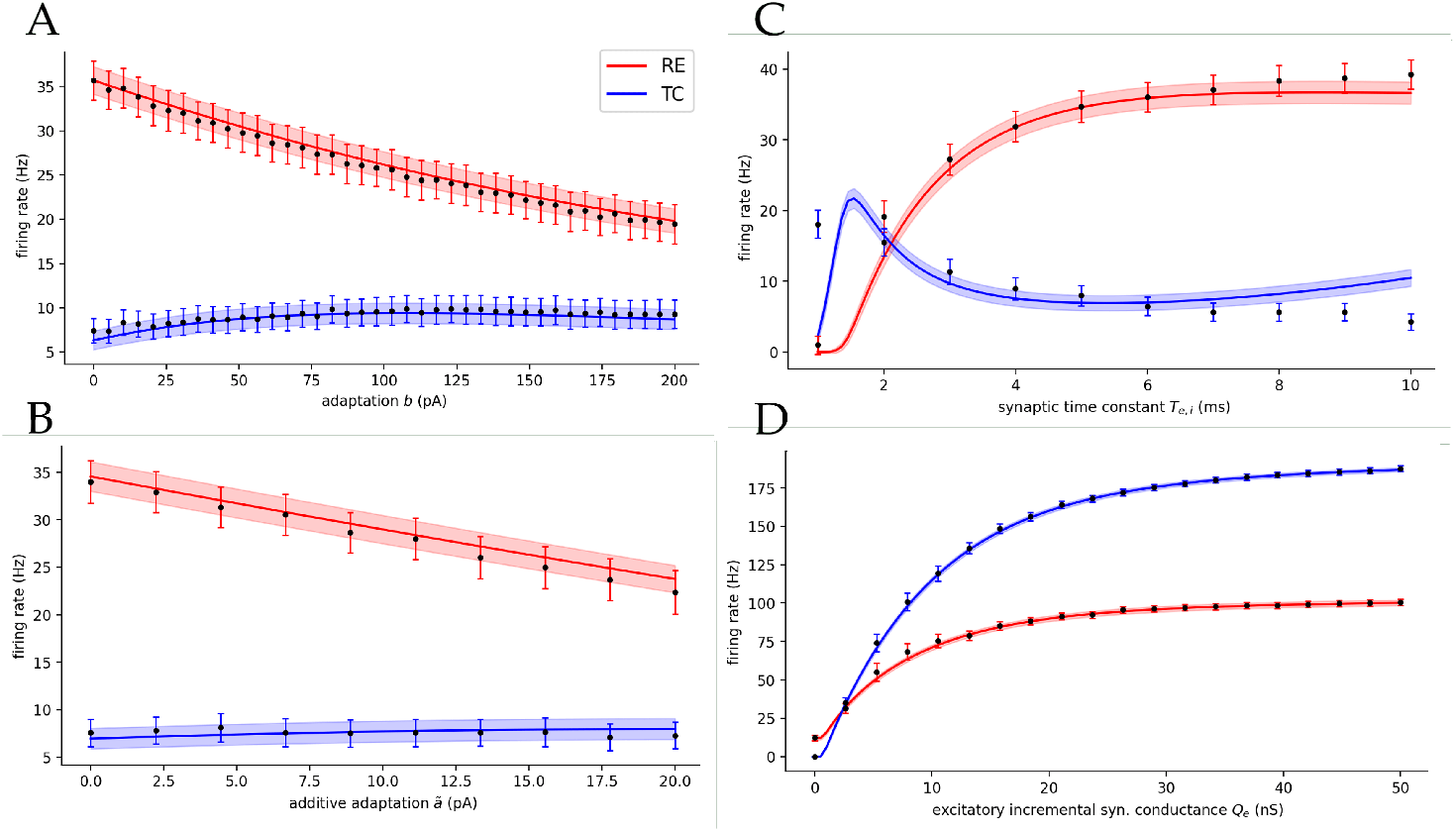
Global parameter analysis for mean-field and spiking network. Black markers represent equilibrium population firing rates of the spiking network. Colored line and shaded area represent the mean-fields mean and standard deviation, respectively. (a) Spiking adaptation parameter. (b) Membrane potential adaptation parameter as shift of the original parameters to keep the parameter. (b) Membrane potential adaptation parameter as shift of the original parameters to keep the difference between TC and RE adaptations, securing stable dynamics. (a) and (b) ensure the mean-fields fit validity between awake and sleep state. (c) Synaptic exponential delay time constant of both populations. (d) Excitatory incremental synaptic conductance *Q*_*e*_ of the TC population (3). The fit good allows for modeling dynamic clamp-like techniques and suggests that the mean-field is able to capture pture the non-trivial firing rate saturation of the spiking network (also supported by Fig. 2d).

**Fig. S4.**
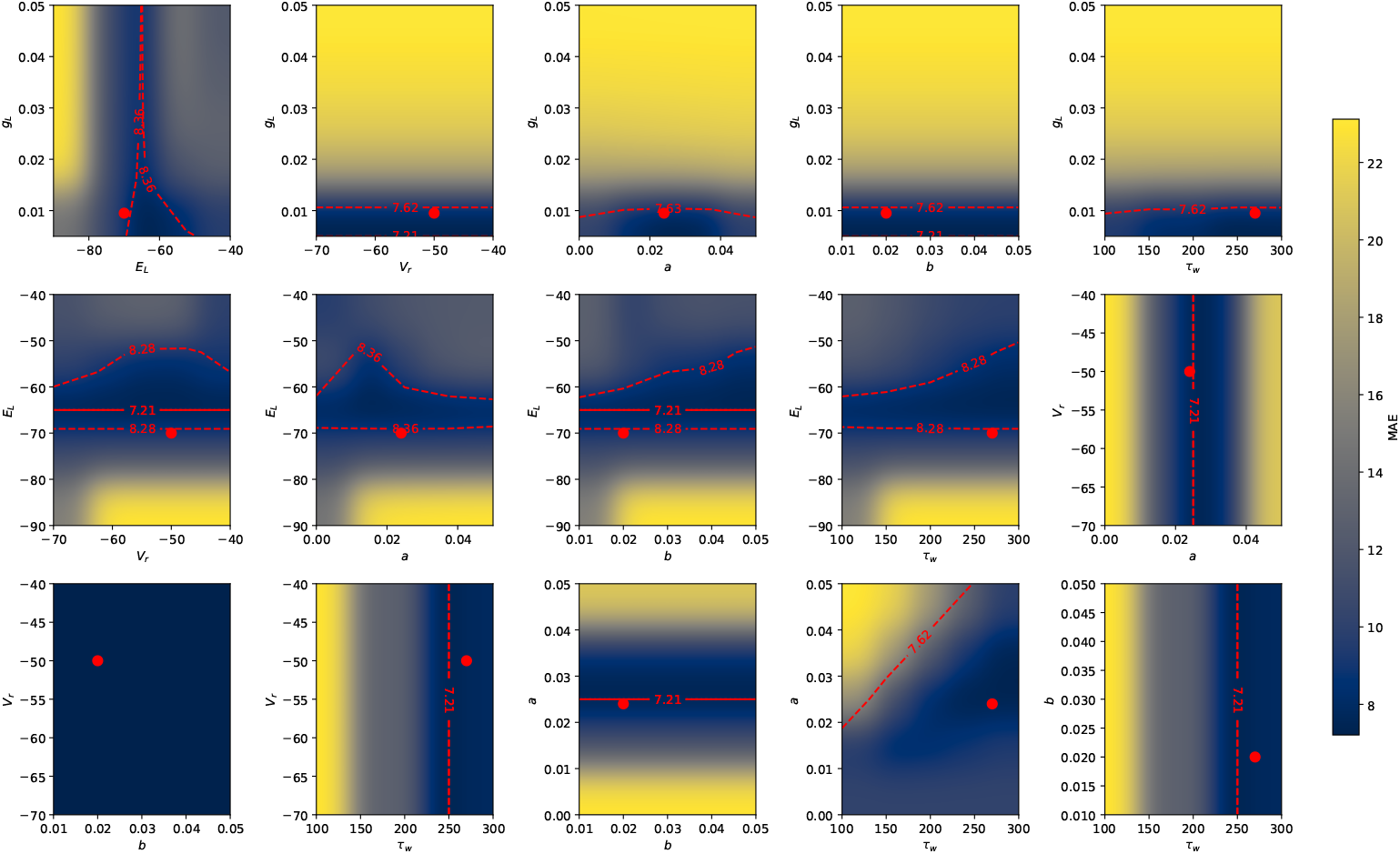
Parameter space fit for TC without ACh. The *mean absolute error* (MAE) between the AdEx (1) and the TC cell traces in absence of ACh from [9]. Red lines mark minimum MAE (solid lines) and 5% deviation from said minimum (dashed lines). The employed parameter values are marked as the red dot. (see units in Fig. S6)

**Fig. S5.**
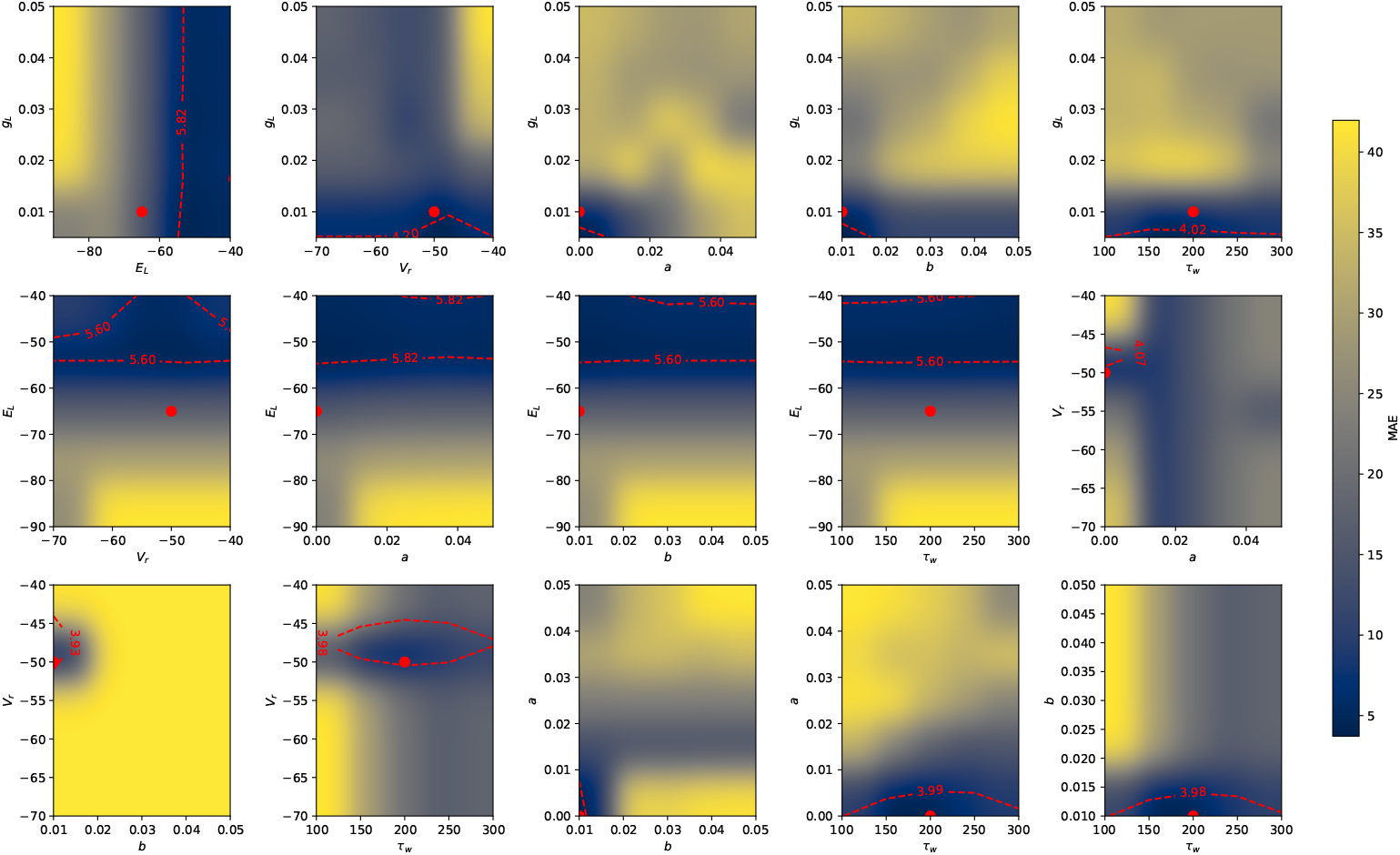
Parameter space fit for TC with ACh. (see caption in Fig. S4)

**Fig. S6.**
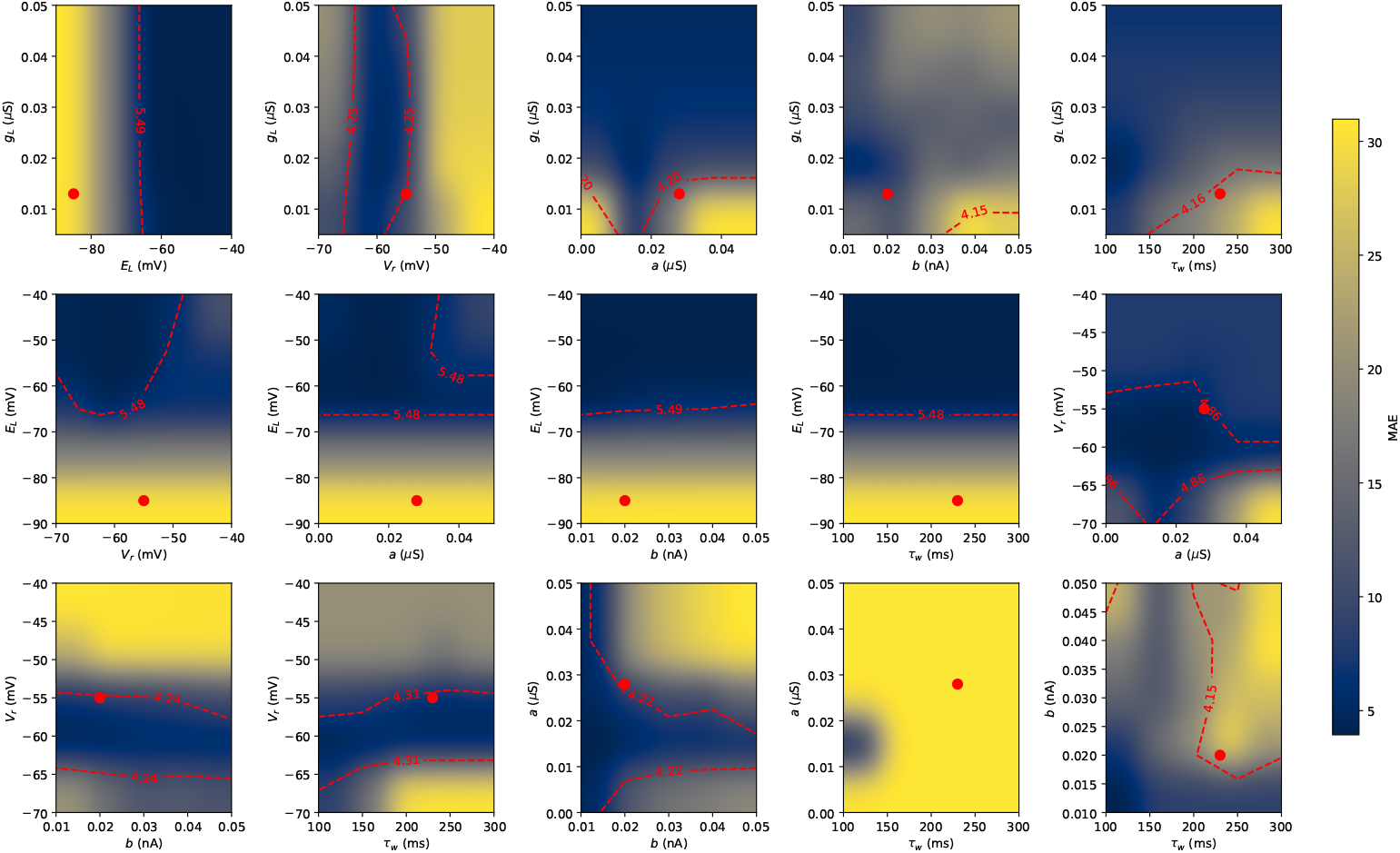
Parameter space fit for RE without ACh. (see caption in Fig. S4) The chosen value of *E*_*L*_ marks the biggest MAE (middle row). Here we still chose to take a hyperpolarised value to account for the reduced excitability in RE neurons with ACh absent [9] and to inhibit also the population response to keep the stability of the network. (see main text)

**Fig. S7.**
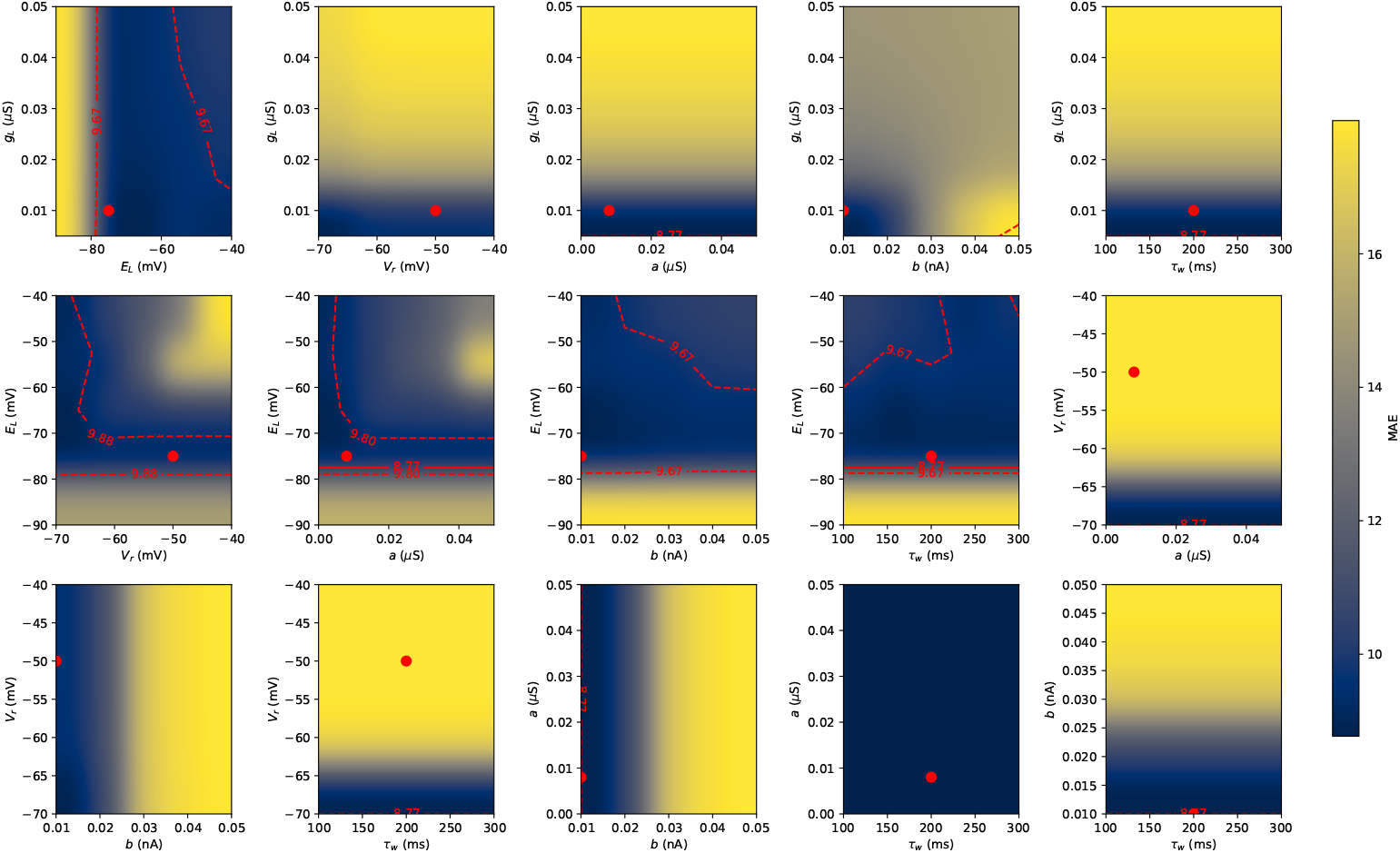
Parameter space fit for RE with ACh. (see caption in Fig. S4)

